# Comparative genetic study of the colony structure and colony spatial distribution between the higher termite *Amitermes parvulus* and the lower, subterranean termite *Reticulitermes flavipes* in an urban environment

**DOI:** 10.1101/2022.12.27.522004

**Authors:** Pierre-André Eyer, Megan N. Moran, Steven Richardson, Phillip T. Shults, Kuan-Ling Kelly Liu, Alexander J. Blumenfeld, Robert Davis, Edward L. Vargo

## Abstract

In insects, ecological competition has often resulted in phenotypic changes and modifications to foraging areas. In termites - and social insects as a whole - colonies cannot easily escape competition through the relocation of their colony. In these species, the outcomes of inter and intra-specific competition are influenced by different life history traits, such as colony size, breeding system (number and types of reproductives), food preference, tunneling patterns, nest site selection, and antagonism between colonies.

Here, we investigated variation in breeding system and spatial distribution among colonies of a higher termite *Amitermes parvulus* and a subterranean termite *Reticulitermes flavipes* within an urban landscape. We first developed microsatellite markers as a tool to study these life history traits in *A. parvulus*. Second, we assessed competitive exclusion or tolerance of *A. parvulus* and *R. flavipes* colonies by determining their fine-scale distribution using monitoring stations on a grid site, and their large-scale distribution across an urban landscape. Third, we investigated the breeding system of *A. parvulus* colonies. We showed that the numerous colonies of *R. flavipes* inhabiting a restricted area contrast with the few, but spatially expansive colonies of *A. parvulus*, suggesting these species face different degrees of intra-specific competition. We showed that colonies of *A. parvulus* frequently merged together, and all of them were headed by inbred neotenic reproductives, two characteristics rarely observed in higher termites. Overall, our study revealed drastic differences in colony structure, breeding systems and foraging ranges between the two species. These differences may reflect differences in food preference and food availability between the two species allowing their co-existence within the same urban environment.

## Introduction

Competition, at both intra- and interspecific levels, directly influences species distribution, diversity, and abundance. It is commonly assumed that multiple species cannot thrive if they overlap within the same, limited ecological niche (*i*.*e*., competitive exclusion theory, Gause 1934; Hardin 1960). Species that are quicker to adapt and respond to environmental changes usually exclude other species through niche substitution and/or competitive exclusion (Byers 2000; Cheng et al. 2009; Holway 1999; McNatty et al. 2009). Although intra- and interspecific competition mostly result in competitors becoming temporally or spatially displaced, they may also lead to physical, physiological and behavioral changes in the life history traits of competing individuals or species that reduce competitive interactions.

Ecological competition within insect groups has resulted in phenotypic changes and modifications to foraging areas (Schoener 1974; Connell 1980). Yet, in contrast to solitary species, the nesting habits of social insect colonies usually prevent these species from easily escaping competition through the relocation of their colony. In these species, the ability to access and secure resources is linked to their ability to exclude competitors through territoriality (Hölldobler & Lumsden 1980). This is particularly true in termites, where colonies of most species are well-defended by a specific soldier caste. It has therefore been suggested that competition represents one of the main regulatory factors controlling termite populations (Noirot 1959; Thorne and Haverty 1991), with the most important competitors of termites being other termites from different colonies or species (Korb and Heinze 2008; Wood and Lee 1971; Perdereau et al. 2011). However, different termite species or colonies may reduce competition by having a broad diet. For example, lower termites have been reported to feed on wet and dry wood, grass and detritus (Eggleton and Tayasu 2001), whereas higher termites, which differ from lower termites by their absence of protozoan symbionts for digesting wood (Eutick et al. 1978; Brune 2014), are known to possess a larger diet range, also including fungus, litter and soil (Eggleton and Tayasu 2001; Donovan et al. 2001). In addition to diet, termite species also exhibit a diversity of foraging strategies, with most species foraging in wood or in the soil using a network of galleries; some higher termites can also forage above ground, harvesting wood debris and plants using trails and/or tunnels (Abensperg-Traun 1993; Miura and Matsumoto 1998; Hu et al. 2012; Almeida et al. 2016). Even among termite species with a common feeding preference and foraging strategy, different species may differ in their nesting and foraging patterns. Some lower termites nest and feed on a single piece of wood (one-piece nesters), while others (both in lower and higher termites) feed on several pieces of wood connected by networks of galleries (multiple-piece nesters), or build large nests separate from their food sources (separate-piece nesters; Abe 1987; Eggleton and Tayasu 2001). Variation in foraging patterns can also be found among species of the same genus (Mizumoto et al. 2020; Janowiecki and Vargo 2022; Pailler et al. 2022), and sometimes even among colonies within species (Mizumoto and Matsuura 2013). Different tunneling patterns have been reported in subterranean termites of the genus *Coptotermes* (Mizumoto et al. 2020), where colonies of *C. formosanus* construct a small number of long tunnels, while *C. gestroi* colonies usually make a large number of short tunnels (Grace et al. 2004, Lee et al. 2007; Hapukotuwa and Grace 2014). Similarly, different drywood termite species of the genus *Cryptotermes* exhibit different tunnel lengths and vary in their foraging preference for different sized pieces of wood (Evans et al. 2011). Overall, these findings suggest that different species or colonies may coexist in a population by possessing differences in their food preferences and foraging habits.

In addition to displaying differences in their foraging habits, termite species also exhibit large flexibility in their colony size and group composition. As termites are social insects, only a few individuals carry out reproduction, whereas the colony duties, such as defense, foraging or brood care, are performed by a large number of non-reproducing workers and soldiers. Foraging behavior is conducted on a colony scale and relies on the collective actions of each individual, mostly workers (Buhl et al. 2005; Traniello and Leuthold 2000). Therefore, distinct features of the group composition may differentially affect termite foraging patterns and strategies. For example, the foraging activity of the workers may vary with the presence and ratio of soldiers (Casarin et al. 2008; Almeida et al. 2016; Janowiecki and Vargo 2022). Similarly, the width and length of the tunnels may differ between those constructed by larger *vs*. smaller workers (Campora and Grace 2004). In addition, foraging patterns can be influenced by both intra- and interspecific interactions, whereby conflicts between colonies or different species can directly result in an alteration of foraging behavior (Traniello 1989; Perdereau et al. 2010; Tian et al. 2017). Furthermore, intra- and interspecific interactions often lead to competitive exclusion in termites when resources are limited (Korb & Linsenmair 2001; Perdereau et al. 2011). Workers of *Reticulitermes flavipes* usually displace other conspecific foragers from a preferred feeding site (Deheer and Vargo 2004) and exclude heterospecific foragers through aggressive behaviors (Perdereau et al. 2011). Finally, larger colonies with a more expansive foraging area would likely outcompete small colonies foraging on a smaller scale. Overall, these findings highlight that colony size, as well as foraging patterns and strategies, directly influence the outcomes of intra- and interspecific competition, facilitating or preventing the coexistence of different colonies within a population (Abe 1987; Reeve and Ratnieks 1993).

Because competitive ability is associated with colony size, different breeding structures may also affect the outcomes of ecological competition by influencing colony growth and capacity. Termites display an extensive variety of breeding structures, even at a small spatial scale (Bulmer et al. 2001; Vargo 2019). Most termite colonies are headed by a monogamous pair of reproductives, a primary king and queen (*i*.*e*., simple families). However, as the colony grows, the death of one or both primary reproductives may trigger the development of worker or nymph offspring into additional reproductives, called neotenics. Although neotenic reproduction is observed in most genera of lower termites (61.7%), it occurs more rarely among genera of higher termites (13.4%, Myles 1999). The presence of multiple functional neotenic reproductives prolongs the life of the colony (*i*.*e*., extended family) and bolsters colony growth, but often results in inbreeding (Bulmer et al. 2001; Vargo and Husseneder 2010). The reproduction of multiple neotenics allows colonies to expand through the environment at a rapid pace (Grube and Forschler 2004). For some species, breeding structure can vary according to changes in environmental factors across a latitudinal gradient (Brandl et al. 2001, 2004; Vargo et al. 2013), or can change throughout time depending upon colony age and interaction with neighboring colonies (Vargo et al. 2003; Vargo 2019; Eyer and Vargo 2022). Although termite colonies may act aggressively when competing with a rival colony over resources, neighboring conspecific termite colonies may occasionally merge, resulting in a cohesive, but genetically diverse colony (*i*.*e*., mixed families, Korb and Roux 2012; Luchetti et al. 2013; DeHeer and Vargo 2004, 2008). The occurrence of mixed families resulting from colony fusion has been mostly observed in lower termites, including dampwood, drywood and subterranean termite species (Goodisman and Crozier 2002; Thorne et al. 2002, 2003; Korb and Schneider 2007; Johns et al. 2009; Vargo and Husseneder 2009). Mixed families have been observed in only a few species of higher termites. Although colony fusion and adoption of unrelated queens have been reported, mixed families of higher termites mostly result from the collaboration of several primary individuals during colony foundation and their persistence throughout the colony’s life cycle (Thorne 1982a, b; Brandl et al. 2001, 2004). These polygynous colonies may dominate a larger spatial area due to a faster growth rate compared to monogynous colonies (Hartke and Baer 2011; Hartke and Rosengaus 2013).

Overall different life history traits, such as colony size, breeding system, foraging strategy, and tunneling patterns, coupled with colony age and antagonism between neighboring colonies, work together to influence competition between conspecific and heterospecific colonies. Because these traits may greatly vary within and between species (especially between a lower and a higher termite species), it suggests that one or more of these factors may be important in determining the spatial distribution of conspecific and heterospecific termite colonies within the landscape. Unfortunately, investigating intra- and interspecific competition in many termites is challenging in the field due to the cryptic nature of their nesting and foraging sites, coupled with the difficulty of identifying species morphologically when only workers are available. Similarly, it is difficult to determine the breeding structure and colony of origin for a collected group of individuals without genetic analyses (Vargo et al. 2003).

Here, we focus on the overlapping distribution between two termite species, the lower termite *Reticulitermes flavipes* and the higher termite *Amitermes parvulus*. To date, the biology of several subterranean *Reticulitermes* termite species is well known, and diverse genetic markers have already been extensively developed and used on these species, especially *R. flavipes* (Dronnet et al. 2005; Perdereau et al. 2013; Janowiecki et al. 2021a; Eyer et al. 2021). Molecular analyses revealed that this species is a multiple-piece nester with native colonies that are often composed of a few satellite nests and feeding sites connected by underground tunnels (usually less than 15 m; DeHeer and Vargo 2004; Shults et al. 2021). These analyses have also highlighted the variability in breeding systems occurring among populations across the eastern US (Vargo et al. 2013). Native *R. flavipes* colonies are usually headed by a monogamous pair of primary reproductives (60-85% of colonies), but they can also be headed by a few neotenics (10-40%), and more rarely they can be the result of colony merging (< 5%) (DeHeer and Vargo 2004; Aguero et al. 2020; Aguero et al. 2021; Vargo 2019; Vargo and Husseneder 2009; Vargo et al. 2013). This indicates that native colonies of this species usually maintain strict boundaries with neighboring colonies.

In contrast to *R. flavipes, Amitermes* is a genus of higher termites (Termitidae). Little research has been conducted to characterize the colony structure and reproductive system of termitid species (Fougeyrollas et al. 2015; Hellemans et al. 2019; Dolejšová et al. 2022), despite the fact that they represent ∼70% of all termite species (Chouvenc et al. 2021). Within the genus *Amitermes*, only the population structure of the ‘magnetic’ termite species *A. meridionalis* from Australia has been genetically investigated (Schmidt et al. 2013). This species is described as a separate-piece nester, with colony centers clearly defined by tall wedge-shaped mounds. A maximum of four alleles per locus has been found within each sampled mound, suggesting that colony foundation is the result of two (unrelated) individuals, and that colony fusion, pleometrosis or the integration of unrelated reproductives (*i*.*e*., mixed families) does not occur. The strong genetic differentiation between mounds suggests that each mound represents a distinct colony. Contrasting with most higher termite species, all colonies of A. meridionalis were extended families (Schmidt et al. 2013). In comparison, no genetic study has investigated the colony structure and reproductive system of *Amitermes* species in the USA, thus no microsatellite markers are available for *A. parvulus*, which has an overlapping geographic distribution with *Reticulitermes* species in central and west Texas. This species does not build mounds; its cryptic subterranean lifestyle therefore hampers accurate determination of colony density and delineation of its foraging strategy.

In this study, we first developed microsatellite markers to detail the breeding system and spatial structure of the termite species *Amitermes parvulus* in central Texas. Second, we assessed competitive exclusion or tolerance of *A. parvulus* and *R. flavipes* colonies by *i*) investigating the fine-scale distribution of the two species using monitoring stations regularly spaced on a grid site, and by *ii*) investigating the large-scale distribution of *Reticulitermes* and *A. parvulus* colonies across an urban landscape. Third, we investigated the colony breeding system of each *A. parvulus* colony sampled by estimating the relatedness among nestmates, the level of inbreeding, and the potential reproduction by neotenics as well as the possibility of colony merging.

## Methods

### Sample collection

#### Assessing fine-scale spatial distribution of colonies within an artificial grid-site

In spring 2016, Advance® Termite Bait Stations (ATBS; BASF) were installed at a field site located in a wooded urban landscape in Pflugerville, Texas, USA with 5 m between each station (*i*.*e*., seven rows of 14 columns; Figure 1a). Each station was made of two wooden monitoring blocks and a blank termite inspection cartridge (*i*.*e*., made of a cellulose bait matrix with no active ingredient; Shults et al. 2021). Stations were first inspected in December 2016, and for each station with termite activity, four auxiliary stations were placed 2.5 m from the main station in each cardinal direction. Stations were re-inspected in December 2017. During this second inspection, approximately 30 individuals were collected and directly stored in 95% ethanol at −20°C for further genetic analyses (Supplementary Information, Table S1) for each station with active termites (both *R. flavipes* and *A. parvulus*).

**Figure 1:**
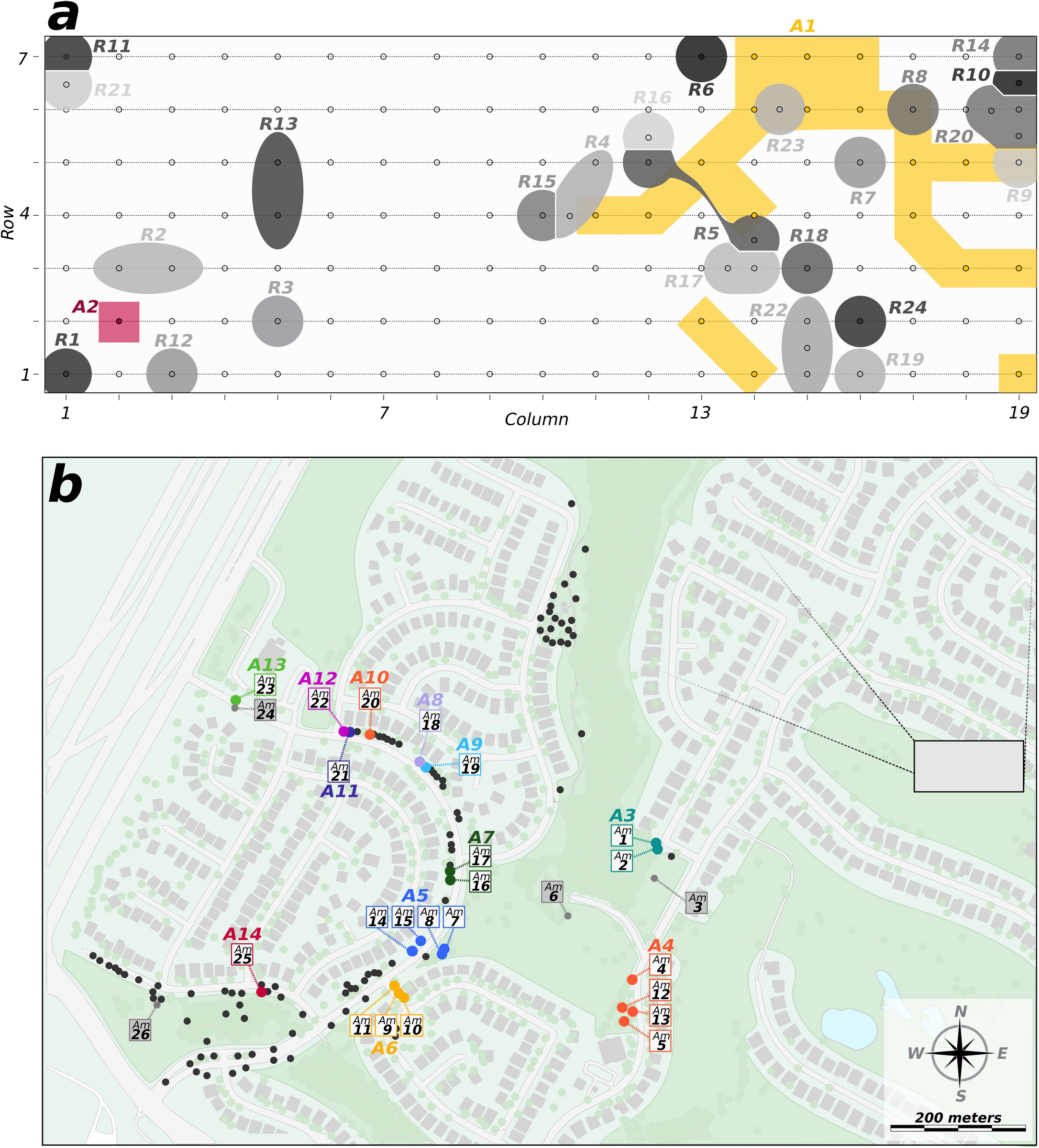
Sampling of *Amitermes parvulus* and *Reticulitermes flavipes* in the fine- and large-scale analyses in the population of Pflugerville, TX, USA. *(a)* In the fine-scale experiment, each small white circle represents an individual Advance station across a grid-site (seven rows x 19 columns). Each active station is illustrated according to the species and colony of origin. Stations of *A. parvulus* are represented by squared shapes and flashy colors (colonies *A1* and *A2*). Stations of *R. flavipes* are represented by rounded shapes and grey-shaded colors (colony *R1* to *R24*). *(b)* In the large-scale experiment, each circle represents a collected sample. Samples of *Reticulitermes* are represented by grey dots. Each sample of *A. parvulus* (*Am1* to *Am26*) is colored according to its colony of origin (colonies *A3* to *A14*); the four grey labels (*Am3, 6, 24* and *26*) represent non-amplifying collection points. The grey inset represents the location of the fine-scale experiment.

#### Assessing large-scale spatial distribution of colonies within an urban landscape

In fall 2021, sampling was conducted within and along trails and streets connecting three parks in Pflugerville, Texas (Mallard Pond Park, Hanging Rock Park, and Secluded Willow Park; Figure 1b). These parks are well-groomed and have a large amount of planting mulch among the trees and vegetation. The top few inches of mulch were removed each time a mulched area was encountered, and every substantial piece of wood was broken open to search for termites. Once termites were encountered, rough species identification was assessed based on soldier morphology (*Reticulitermes* vs. *Amitermes*) and geospatial coordinates were recorded. *Reticulitermes* samples were discarded, and *Amitermes* samples were taken back to the lab for further genetic analyses.

### Molecular and genetic analyses

A total of 34 *R. flavipes* and 21 *A. parvulus* samples were collected for the fine-scale analysis, and an additional set of 26 *A. parvulus* samples were collected for the large-scale analysis from the three parks in Pflugerville (98 points of *R. flavipes* were encountered within the parks, but not collected). From each collected sample, four to 10 individuals were genotyped to infer colony assignment (*R. flavipes* and *A. parvulus* samples) and breeding system (*A. parvulus* samples only).

Total genomic DNA was extracted from the head of each individual using a modified Gentra Puregene extraction method (Gentra Systems, Inc. Minneapolis, MN, USA). Note that a high number of non-amplifications was obtained when extracting entire *A. parvulus* termites during preliminary testing. In addition, DNA was extracted from a pool of heads from 50 *A. parvulus* individuals to ensure high DNA quantity; this sample was sent to Novagene for Illumina sequencing on a NovaSeq S2 to identify microsatellite repeats.

### Microsatellite primer design and amplification

After Trimmomatic v. 0.39 was used to filter the raw sequenced reads for quality (SLIDINGWINDOW:4:25) and Illumina adapter contamination (ILLUMINACLIP; Bolger et al. 2014), ABySS v. 2.1.5 was used to assemble the cleaned reads into contigs for subsequent microsatellite identification (Jackman et al. 2017). Finally, the software QDD v. 3.1 (Meglécz et al. 2014) was used to discover microsatellite repeat motifs within the cleaned, assembled reads derived from the pool of *A. parvulus* heads. A threshold of at least five repetitions, excluding mononucleotide repeats, was established for subsequent primer design. Overall, 1,259,044 reads containing microsatellite repeat motifs were identified. For each of those reads, the 200 bp flanking regions on either side of the repeats were extracted. Microsatellite motifs with the highest number of repeats were chosen to maximize polymorphism. We selected a set of 25 loci and designed the corresponding primers within both flanking regions using the online Primer3 software (http://primer3.ut.ee). These primers were designed to result in a broad range of PCR product sizes to facilitate multiplex arrangements.

For *A. parvulus*, the 25 primer pairs were first tested in standard simplex PCR conditions on a small number of individuals. Markers were discarded if they were monomorphic or did not amplify consistently. Preliminary testing allowed us to select a set of 16 microsatellite markers that amplified and were polymorphic in this preliminary set of samples. Primer sequences, microsatellite repeat information, PCR conditions, multiplex arrangements, and allele numbers for those 16 microsatellite markers are provided in Table 1. The M13-tailed primer method was implemented to allow for multiplexing. The M13 tails were attached to the forward primer and 5’-fluorescently labeled with 6-FAM, VIC, PET, or NED dyes. PCR was performed using a Bio-Rad thermocycler T100 (Bio-Rad, Pleasanton, CA, USA). Prior to genetic analyses, the software Genepop v.4.7.0 (Rousset 2008) was used to verify the absence of linkage disequilibrium between each pair of newly designed microsatellite markers (All *P* > 0.05). For *R. flavipes*, two highly polymorphic and unlinked microsatellite markers (*Rf21-1* and *Rf24-2*) were amplified for each individual of the finescale analysis following the method of Vargo (2000). The resulting products for both species were visualized on an ABI 3500 capillary sequencer and sized against a LIZ500 internal standard (Applied Biosystems, Foster City, CA, USA). Geneious v.9.1 (Biomatters, Auckland, New Zealand; Kearse et al. 2012) was used to score alleles.

**Table 1:**
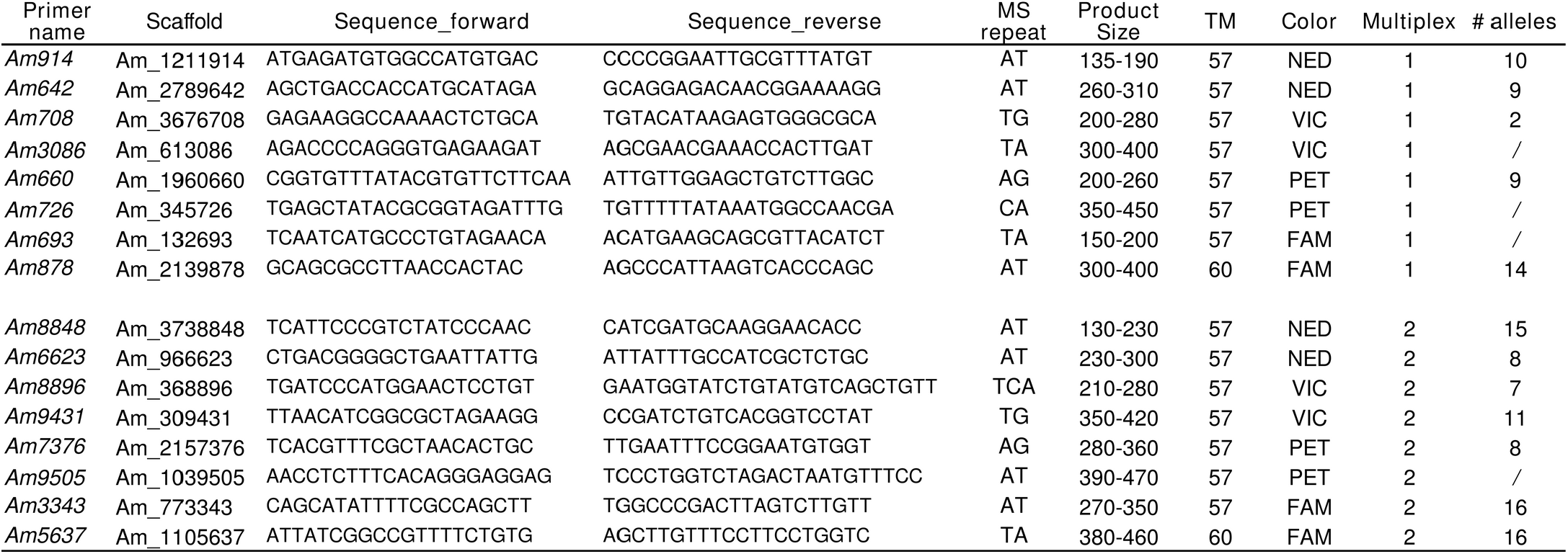
Primer sequences, microsatellite repeat information, product size, PCR condition, multiplexing arrangement, and allele number for each microsatellite marker.

### Genetic assessment of colony assignment

A total of 329 individuals were genotyped to determine the colony identity for the 34 *R. flavipes* samples from the grid-site stations. For *A. parvulus*, 155 and 143 individuals from the 21 and 26 samples were genotyped to identify colony boundaries for the fine- and large-scale analyses, respectively (Table S1).

The genotypes of workers between each pair of samples were used to identify whether workers from different collection samples belong to the same or distinct colonies. Pairwise *F*_*ST*_ values were calculated, and genotypic differentiation was tested for every pair of samples using a loglikelihood (G)-based test in Genepop v.4.7.0 (Rousset 2008). Two samples were considered to belong to the same colony when they exhibited a low pairwise *F*_*ST*_ value and when allelic frequencies did not differ significantly based on the Gtest (*P* < 0.05 after Bonferroni correction). For *A. parvulus*, the assignment of samples into distinct colonies was also visualized using Bayesian and PCA clustering approaches. Bayesian clustering was implemented in Structure v.2.3.4 (Pritchard et al. 2000). For each analysis, simulations were run with K ranging from one to the number of samples, with 10 replications for each value of K. Each run comprised of a first step with a 50,000 burn-in period and 100,000 iterations of the Markov chain Monte Carlo (MCMC) simulation algorithm. The method based on the variation of LnP(k) (Pritchard et al. 2000), implemented in StructureSelector (Li and Liu 2018), was used to determine the most likely number of genetic clusters. Then, individual genotypes were examined using a principal component analysis (PCA) implemented in the *adegenet* R package (Jombart 2008). Workers from samples belonging to the same colony should cluster together, while workers from different colonies should segregate across the axes. Because the samples of the finescale analysis only belong to two distinct *A. parvulus* colonies (see Results), *F*-statistics, as well as Bayesian and PCA clustering analyses were performed together for the fine-scale and large-scale analyses to better estimate the allelic frequencies within the overall population.

Species and colony identities were further confirmed through sequencing of the mitochondrial cytochrome oxidase II gene for one individual for each *A. parvulus* sample using *A-tLeu* and *B-tLys* primers. A haplotype network was used to visualize genetic divergence between mitochondrial haplotypes. For simple- or extended-family colonies, individuals from collection samples belonging to the same colony are expected to carry the same mitochondrial haplotype. Networks were produced by the median-joining method implemented in the program Network v.4.6.1.1 (Bandelt et al. 1999).

### Genetic determination of *A. parvulus* breeding system

For breeding system analyses, all the collected samples of *A. parvulus* belonging to the same colony, based on the analyses mentioned above, were combined. First, relatedness coefficients (*r*) between workers were estimated for each colony using Coancestry v.1.0 (Wang 2011), following the algorithm described by Queller and Goodnight (1989). The family type of each colony (*i*.*e*., simple-, extended- or mixed-family) was identified based on the number and frequency of alleles within each colony. Colonies exhibiting more than four alleles at one or more loci were classified as mixed families. When no more than four alleles at any locus were observed, colonies were categorized as *i*) an extended family when genotypic combinations were inconsistent with a monogamous pair of reproductives (*e*.*g*., an allele pairing with itself and two others) and when high level of *F*_*IS*_ was observed (*F*_*IS*_ > 0), or *ii*) a simple family (Vargo 2003). The inbreeding coefficient *F*_*IS*_ and observed heterozygosity were calculated for each colony using the software Fstat (Goudet 1995).

In addition, to determine if there was a gradual increase in genetic differentiation among colonies with geographical distance due to limited dispersal, the genetic isolation by distance was tested among *A. parvulus* colonies. The genetic differentiation coefficients [*F*_*ST*_ /(1 – *F*_*ST*_)] between pairs of colonies were plotted against the *ln* of their geographical distances (Slatkin 1993). The Mantel test implemented in Genepop (Rousset 2008) was used to test the significance of the correlation.

## Results

Among the 16 microsatellite markers selected from preliminary analyses of *A. parvulus*, 12 markers amplified consistently and yielded clearly discernible results in most individuals; the four other markers were later removed due to nonamplification on a substantial number of samples. The 12 remaining microsatellite markers revealed a number of alleles ranging from two to 16 (*mean* ± *SD* = 10.42 ± 4.21; Table 1) and can therefore be used as an informative tool for analyzing the genetic structure of *A. parvulus* colonies. Because this study only includes individuals originating from a single population, these markers are likely to exhibit higher polymorphism across the entire distribution range of the species; therefore, they can also be used to provide robust insights into the population genetics of this species at larger geographic scales.

### Fine-scale spatial analysis

Soldier-morphology-based species identification revealed that both *R. flavipes* and *A. parvulus* were found among stations in the fine-scale analysis. Preliminary genotyping revealed that two stations were simultaneously occupied by both species in different parts of the station; for these stations, eight individuals were genotyped for each species.

The markers *Rf21-1* and *Rf24-2* were highly polymorphic in *R. flavipes* (31 and 30 alleles respectively), and thus sufficient to assess colony identification, as previously done by Shults et al. (2021) in a similar sampling-designed experiment. Over the 32 active stations with *R. flavipes*, 24 distinct colonies were identified using the G-test of genotypic differentiation (Figure 1a). Overall, the mean pairwise *F*_*ST*_ values between collection samples belonging to the same colony was 0.012 ± 0.041 (± *SD*; ranged from -0.039 to 0.103), while the mean pairwise *F*_*ST*_ values between collection samples belonging to different colonies was 0.277 ± 0.065 (± *SD*; ranged from 0.072 to 0.453). The pairwise *F*_*ST*_ values between every pair of collection samples are provided in Supplementary Material (Figure S1).

Over the 21 active stations with *A. parvulus*, only two distinct colonies were identified using the G-test of genotypic differentiation (Figure 1a). Therefore, the mean number of active stations was similar between the two species; however, it resulted in a drastic difference in the number of colonies. Consequently, the mean number of active stations per colony for *R. flavipes* was 1.41 ± 0.72 (± *SD*; ranged from 1 to 3) and up to 20 active stations for one colony of *A. parvulus*. Although the only other colony was found within a single active station, we cannot rule out that this second *A. parvulus* colony further extended outside of the studied area. As a result, the largest distance between two *R. flavipes* sampling stations belonging to the same colony was 12.5 m (two columns and 1.5 rows, Colony *R5*), while it was 40.3 m (eight columns and 1 row) between the two furthest sampling stations belonging to the same *A. parvulus* colony (Colony *A1*). It also resulted in the closest distance between two *R. flavipes* colonies being 2.5 m (0.5 station), while it was 46.1 m (nine columns and 2 rows) between the two *A. parvulus* colonies (Figure 1a).

### Large-scale spatial analysis

Of the 119 encounters with termites across the urban landscape in central Texas, 82% belonged to *Reticulitermes* (N = 98, Figure 1b). Although we did not genetically determine the colony identity of these collection samples, the average distance between two collection samples of the same *R. flavipes* colony and the shortest distance between two *R. flavipes* colonies observed in the fine-scale analysis suggest these 98 sample collections potentially represent between 79 (19 pairs of collection points were located less than 12.5 m away from each other) and 98 (all collection points are located more than five meters away from each other) different colonies.

Four of the 26 collection points of *A. parvulus* did not amplify using the set of newly designed microsatellite markers (*Am 3, 6, 24, 26*), probably as a result of poor DNA preservation or the presence of a biological compound within individuals preventing DNA amplification. The 22 remaining collection points were genetically assigned to 12 distinct colonies of *A. parvulus* (Figure 1b). Similar to the results revealed in the finescale analysis, the largest distance between two collection points belonging to the same *A. parvulus* colony was 56.8 m, with an average distance of 25.8 m ± 18.3 (± *SD*; ranged from 1.5 to 56.8). If we consider the four *A. parvulus* collection points that did not amplify as distinct colonies, the average distance between one *A. parvulus* colony to the closest conspecific colony was 65.8 m ± 59.5 (± *SD*; ranged from 5.7 to 161.3 m).

Based on both fine- and large-scale analyses, the mean pairwise *F*_*ST*_ value between collection samples belonging to the same *A. parvulus* colony was 0.04 ± 0.05 (± *SD*; ranged from -0.04 to 0.21), while it was 0.43 ± 0.08 (± *SD*; ranged from 0.19 to 0.75) between samples from different colonies. The pairwise *F*_*ST*_ values between *A. parvulus* collection samples are provided in the Supplementary Material (Figure S2). The genetic clustering of the 21 and 22 collection points into the two and 12 genetically inferred colonies of the fine- and large-scale analyses, respectively, was also evident in the PCA, as the different colonies scattered along the axes, with all individuals/collection points from a given colony segregating together (Figure 2a, b). Structure analysis suggested a similar finding with an optimal number of genetic clusters also being 11 based on the LnP(k) method (Figure 2c). At this value of K, most individuals within the 11 genetic groups have a high probability of assignment to their respective cluster (*i*.*e*., colony). However, in contrast with the analyses above, some pairs of colonies could not be genetically distinguished from each other (Colonies *A4* and *A8*; *A3* and *A13*, as well as *A9* and *A10*). However, it is unlikely these pairs of sampling points belong to the same colony, as they are geographically distant from each other (respectively 476m, 660m, and 98m; Figure 1b). Unfortunately, the sequencing of the mitochondrial marker was not informative in distinguishing colonies, as the population was comprised of only three haplotypes, one of them being shared by 90% of the sample points (Figure 3a). Therefore, this sequencing analysis can only confirm the separation between colonies *A4* and *A8* (*i*.*e*., carrying different haplotypes), which was unclear based on the results of Structure but was unlikely due to their geographic distance.

**Figure 2:**
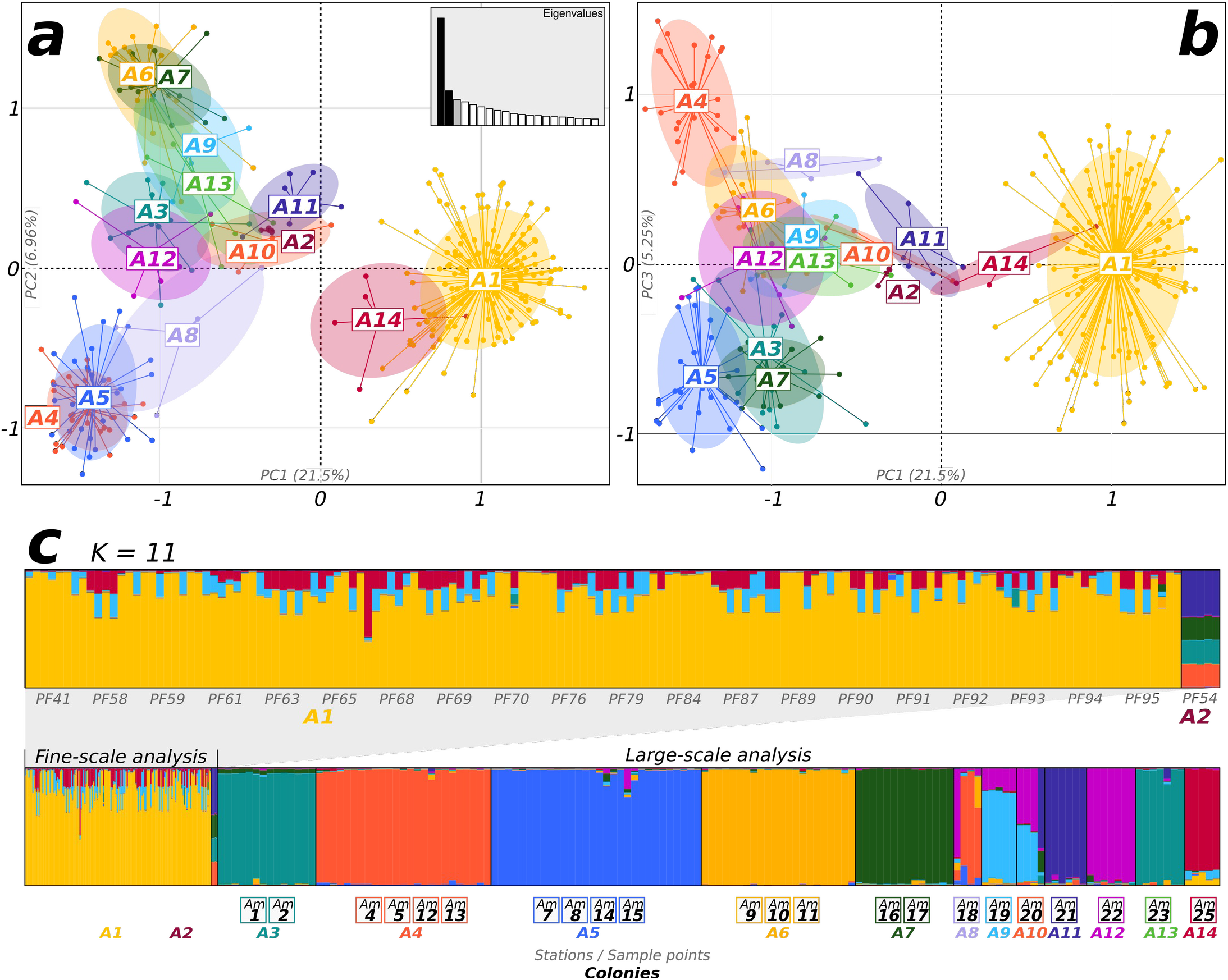
*(a, b)* Principal Component Analysis (PCA) of the microsatellite markers for all the collection points of *A. parvulus* sampled from both the fine- and large-scale analyses. The first and second axes of the PCA are represented on the left panel *(a)*; the first and third axes are represented on the right panel *(b)*. Each individual is colored according to its colony of origin (colonies *A1* to *A14*). *(c)* Graphical representation of Structure results for K = 11 genetic groups in the overall *A. parvulus* dataset (*i*.*e*., fine- and large-scale analyses). Each group is characterized by a color. Each individual is represented by a vertical bar colored according to its probability of assignment to each group. The first plot (above) only included the individuals of the fine-scale analysis. This plot is only a zoom in of the overall plot and does not correspond to a separate Structure analysis.

**Figure 3:**
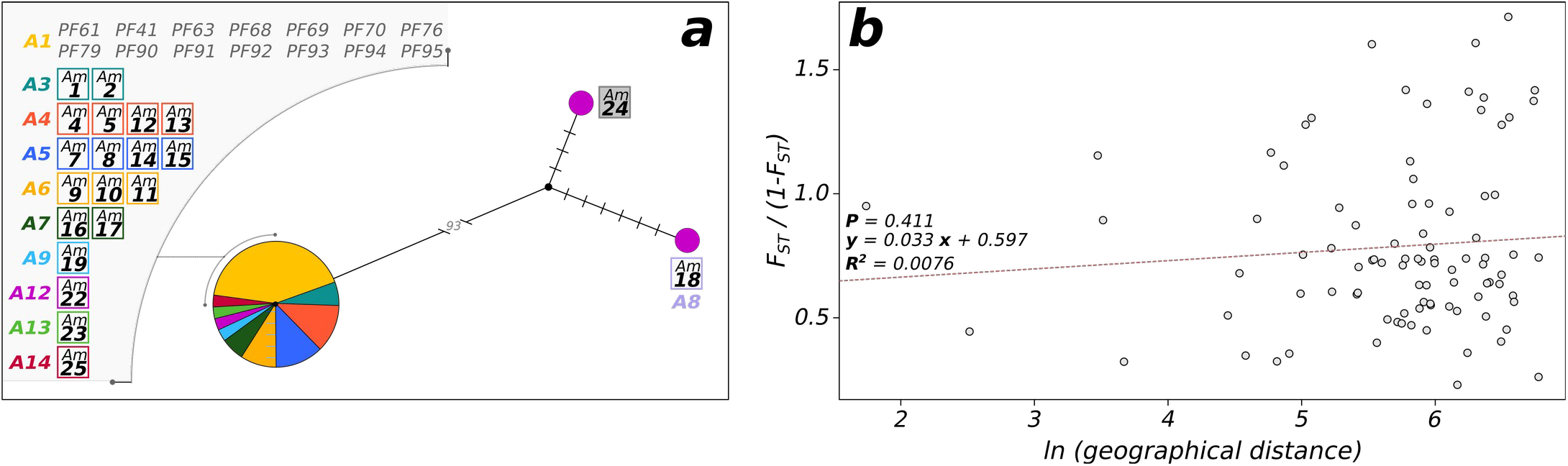
*(a)* Haplotype network for the COII mitochondrial marker for one individual for each of the *A. parvulus* collection samples. Circle sizes are proportional to the number of sequences observed in the dataset. Each sample is colored according to its colony of origin (colonies *A1* to *A14*). Branch lengths and tick marks indicate the number of mutations between haplotypes. (b) Correlations of genetic differentiation between colonies [*F*_*ST*_ /(1 *F*_*ST*_)] and the ln of their geographic distances. This analysis uses the different colonies of *A. parvulus* (*A1* to *A14*), but not the different collection points (*Am1* to *Am26*).

### Breeding system analyses

The 21 and 22 collection samples of *A. parvulus* were combined into two and 12 colonies from the fine and large-scale analyses, respectively. Among these 14 *A. parvulus* colonies, the majority were extended families (N= 9) and mixed families (N = 4), with only a single simple family found (Table 2). Notably, the unique simple family was inferred based on its highly negative level of inbreeding (*F*_*IS*_ = -0.467), but the presence of only 2 alleles for each of the markers could also correspond to the distribution of genotypes expected under an extended family. Overall, the mean level of inbreeding within each colony (*F*_*IS*_) was 0.109 ± 0.230 (± *SD*). The mean level of inbreeding was 0.118 ± 0.189 (± *SD*; ranged from –0.125 to 0.510) within extended families, and 0.230 ± 0.055 (± *SD*; ranged from 0.169 to 0.300) within mixed families. This high level of inbreeding resulted in the observed heterozygosity (*H*_*o*_ = 0.373) being lower than the expected heterozygosity (*H*_*e*_ = 0.780). Interestingly, the high level of inbreeding also found in mixed-family colonies suggests those colonies arose from the merging of two extended families (Thorne et al. 1999). Overall, the common occurrence of inbreeding, even in mixed families, resulted in a high level of relatedness among workers within colonies, with an average relatedness of 0.554 ± 0.164 (± *SD*; Table 2). The mean relatedness was 0.576 ± 0.129 (± *SD*; ranged from 0.408 to 0.796) within extended families, 0.419 ± 0.089 (± *SD*; ranged from 0.332 to 0.538) within mixed families, and 0.893 among workers of the simple family. The Mantel test indicated an absence of significant correlation when examining the genetic differentiation between pairs of colonies and their geographic distances (*P* = 0.411; Figure 3b), suggesting a large genetic mixing across the urban landscape studied.

**Table 2:** Sample locations, family type, inbreeding coefficient and relatedness for each genetically inferred colony of *A. parvulus*.

| Analyses | Colony ID | Species | Stations / sampling points | # stations | Family Type |
| --- | --- | --- | --- | --- | --- |
| Fine-scale | A1 | *Amitermes* | all but PF54 | 20 | Mixed |
| Fine-scale | A2 | *Amitermes* | PF54 | 1 | Simple |
| Large-scale | A3 | *Amitermes* | Am1, 2 | 2 | Mixed |
| Large-scale | A4 | *Amitermes* | Am4, 5, 12, 13 | 4 | Mixed |
| Large-scale | A5 | *Amitermes* | Am7, 8, 14, 15 | 4 | Extended |
| Large-scale | A6 | *Amitermes* | Am9, 10, 11 | 3 | Extended |
| Large-scale | A7 | *Amitermes* | Am17, 16 | 2 | Extended |
| Large-scale | A8 | *Amitermes* | Am18 | 1 | Extended |
| Large-scale | A9 | *Amitermes* | Am19 | 1 | Extended |
| Large-scale | A10 | *Amitermes* | Am20 | 1 | Extended |
| Large-scale | A11 | *Amitermes* | Am21 | 1 | Extended |
| Large-scale | A12 | *Amitermes* | Am22 | 1 | Mixed |
| Large-scale | A13 | *Amitermes* | Am23 | 1 | Extended |
| Large-scale | A14 | *Amitermes* | Am25 | 1 | Extended |

## Discussion

Our study provides a set of microsatellite markers enabling the empirical investigation of *Amitermes parvulus* colony structure and population genetics in the US. In this study, the newly developed microsatellite markers were used to investigate the colony distribution of this species within a grid-site experiment and across an urban landscape. By using a direct comparison between the colony distribution of *A. parvulus* and *Reticulitermes flavipes* on the same grid site, our study provides a better understanding of the foraging activities and colony distribution of these two termite species in Central Texas. Our study also reveals that a substantial proportion of colonies of *A. parvulus* were mixed families, and all of them were headed by inbred neotenic reproductives, two characteristics rarely observed in higher termites. This provides insights into both intra- and interspecific competitive pressures acting in these populations. Our study revealed drastic differences between the two species in terms of breeding systems, the size of their foraging range, and the size of their foraging buffer with their neighboring conspecific colonies. These differences probably reflect their differences in habitat use (open areas *vs*. forests) and in food preference (small twigs *vs*. large pieces of wood) that may potentially reduce their niche overlap, allowing them to coexist within the same locality.

For *R. flavipes*, our study revealed that colonies inhabited a limited foraging area, with a maximum of three stations per colony. Consequently, the 24 colonies uncovered across the grid site roughly correspond to 90 colonies per hectare. Similar results were previously reported for this species using a similar experimental design in two other Texan populations (located∼120 and 200km away), identifying few stations per colony and 55-120 colonies per hectare (Janowiecki and Vargo 2021b; Shults et al. 2021). Our fine-scale analysis found that the maximum distance between two collection points of *R. flavipes* belonging to the same colony was ∼12.5 m; these results match those reported for field experiments indicating that two collection points located more than 15 m away from each other probably belong to two different colonies (Vargo 2003; DeHeer and Vargo 2004). In contrast, no previous results on the geographic distribution of colonies were available for *A. parvulus*, or any *Amitermes* species in the Nearctic region. The colonies of *A. parvulus* seem to be spatially larger (up to four times) than those of *R. flavipes*. Consequently, both our fine- and large-scale analyses revealed that the distance between two distinct colonies of *A. parvulus* is higher than the distance between two *R. flavipes* colonies. The greater distances between *A. parvulus* colonies may suggest high intraspecific competition between colonies of this species or may simply reflect the low quantity of nesting habitats available in the studied population. Species of the genus *Amitermes* are commonly found in arid areas, feeding on small twigs and grass stems. For our large-scale analysis, all apparent wood material was investigated, such as mulch and pieces of broken branches. Although a few unsuccessful collection points in grassy areas were performed, we did not fully investigate the presence of termite colonies under large grassy areas, feeding on grass stems, root material or potential buried pieces of wood. However, the presence of the two species within the grid site of the small-scale analysis (*i*.*e*., made of wooden monitors buried mainly in open pastures) suggests their presence in such areas. For both species, the potential presence of additional colonies in these unsampled areas may reduce the mean distance between two colonies. Although the colonies of *A. parvulus* are more spatially expansive than those of *R. flavipes*, we do not know whether *A. parvulus* colonies are more populous than those of *R. flavipes* or whether they simply extend over a larger foraging range. Similarly, we do not know whether the multiple sample points collected for each *A. parvulus* colony represent foraging groups of workers outside their central nest (separate-piece nesters) or multiple satellite nests (multiple-piece nesters). Interestingly, some collection points of this study consisted of large pieces of wood opened in wooden areas/forests within the parks, in which only dense nests of *R. flavipes* were found. These results suggest that *A. parvulus* probably does not forage for large pieces of wood, potentially lowering interspecific competition with *R. flavipes*, but instead forages underground over a spatially extensive range in open areas. Colonies of the congeneric species *A. wheeleri* cannot detect food below a depth of 15-20cm; instead, they mostly forage near the surface of the soil, locating food items above the soil surface by sensing their thermal shadows (Ettershank et al. 1980). This foraging strategy was also reported in the desert species of the sister genus, *Gnathamitermes tubiformans*, foraging underground in a similar habitat. Overall, these findings suggest that, in contrast to *R. flavipes, A. parvulus* probably does not thrive in large, covered areas, such as forests. In addition, the difference in the size of the consumed food source (small twigs *vs*. large pieces of wood) may also explain the difference in foraging ranges between the two species, where colonies of *A. parvulus* need a larger foraging area to acquire a similar quantity of resources.

In higher termites, the presence of mixed families has been observed in only two species, *Macrotermes michaelseni* and *Nasutitermes corniger* (Brandl et al. 2001; Thorne 1984), mostly through pleometric association of co-founding primary individuals. Our results add to the list of known mixed families in higher termites -with a substantial proportion of colonies (∼28%) being mixed families-although we do not know whether they originate from colony fusion, pleometrosis, or both. However, our results for *A. parvulus* differ from the findings for *A. meridionalis*, the only other species of *Amitermes* for which colony structure was genetically studied, in which none of the 99 colonies investigated were mixed families (Schmidt et al. 2013). Although, the number of reproducing neotenics was not investigated in this study, our analyses reveal the reproduction by neotenics in all colonies investigated, suggesting that workers of this species may readily develop into neotenic reproductives. Although neotenic reproduction is relatively uncommon among genera of higher termites, it is more common among the *Amitermes* group than in any other Termitidae, with some species (*e*.*g*., *A. laurensis*) having more neotenic headed colonies observed than colonies headed by primary reproductives (Myles 1999). Similarly, in *A. meridionalis*, all colonies investigated were headed by neotenic reproductives, leading to a high level of inbreeding within colonies (Gay & Calaby 1970; Schmidt et al. 2013). Coupled with our results, these findings suggest the widespread occurrence of extended families among the species of the genus *Amitermes*. To date, the colony genetic structures of higher termite species have been barely investigated even though they represent most termite species worldwide. Our results highlight the variability of colony structure in this group, even within a given genus, calling for further investigation of this life history trait for more termitid species.

*Reticulitermes flavipes* is a successful invasive species, introduced to many regions worldwide, such as France, Chile, and Uruguay (Clément et al. 2001; Austin et al. 2005; Su et al. 2006; Eyer et al. 2021), while *A. parvulus* has not been reported as introduced outside of its endemic range. The spatially expansive foraging range of *A. parvulus* is a potential characteristic enhancing invasiveness, as it increases the chance of being inadvertently picked up and transported abroad (Evans et al. 2011, 2013; Pailler et al. 2022); however, this species was mostly found tunneling in mulch areas but not inhabiting substantial pieces of wood. The absence of dense colony fragments within large pieces of wood may have restricted the opportunity for large fragments of colonies to be transported. Additionally, the invasion success of *R. flavipes* has been, at least partially, attributed to its breeding system. Colonies of the French invasive range are usually large, headed by several hundred neotenic reproductives and readily merge together (Dronnet et al. 2005; Perdereau et al. 2010; Vargo and Husseneder 2009). As mentioned above, colonies of *R. flavipes* in the native range usually do not share these characteristics, except for a population from Louisiana that has been suggested as the potential source population of the invasion (Perdereau et al. 2010, 2015); yet this population may have arisen through a reintroduction from France to Louisiana (Eyer et al. 2021). In contrast, colonies of the non-invasive species *A. parvulus* are similar to the invasive colonies of *R. flavipes*, as they are spatially expansive, a substantial proportion of them are mixed families, and all of them are headed by neotenic reproductives. The swift development of neotenic reproductives from nymphs or workers is also known to favor invasiveness as it allows every group of transported workers to develop into a viable propagule (Evans et al. 2011, 2013; Eyer and Vargo 2021). However, despite being one of the most destructive invasive termite species, most mature colonies of *C. formosanus* are headed by primary reproductives in its introduced range in the US (Husseneder et al. 2005, 2012; Blumenfeld et al. 2021). Overall, these findings further suggest that breeding systems may not be as strong of a factor when it comes to influencing invasiveness in termites as it is in ants (McGlynn 1999; Holway et al. 2002; Eyer and Vargo 2021), indicating that other factors or opportunities may better explain the likelihood of a termite species to become established outside of its native range (Blumenfeld and Vargo 2020). These results also highlight the need for expanding our basic knowledge about colony structure and breeding systems for more termite species, including non-pest or non-invasive species, to draw robust conclusions about life-history traits influencing invasiveness in termites.

The apparent overlap between colonies of *A. parvulus* and colonies of *R. flavipes*, sometimes inhabiting the same stations (Figure 1a), calls into question whether *A. parvulus* exhibits highly aggressive behavior toward heterospecific termite species, such as *R. flavipes*, and how the winner of these confrontations may be determined. In addition, the high occurrence of mixed families in *A. parvulus* also raises the question of how distinct colonies of *A. parvulus* interact when they come into contact with each other, therefore calling for the use of behavioral assays to test for intraspecific aggression or merging (Leponce et al. 1996; Korb and Roux 2012; Cooney et al. 2016). Figure 1b shows a fairly even separation of colonies (possibly indicative of territoriality), though this can be achieved by factors other than aggression, such as avoidance (Evans et al. 2009; Li et al. 2010). Our data also revealed that colonies of this species are spatially expansive, with some foraging galleries or satellite nests disconnected from the main nesting area (Figure 1a). These foraging characteristics may increase colony merging between disconnected fragments of colonies; however, this can only occur if aggression between these fragments is low. Interestingly, these findings may also explain the increase of worker development into neotenic reproductives within outer satellite nests (Myles 1999). Excavation of colonies will help to determine the number of neotenics within colonies, as well as whether they are gathered in the center of the colony or spread across numerous satellite nests, as these characteristics are important for understanding colony dynamics and growth. In addition, it would be interesting to investigate neotenic reproduction within merged colonies to determine whether only the neotenics of a single original colony keep reproducing, whether they interbreed, and/or whether colony merging creates opportunities for nonrelative workers to preferentially develop into neotenics (Johns et al. 2009).

The absence of isolation by distance across the population indicates that colony spread still arises through the nuptial flight of primary reproductives. This dispersal mode allows founders to establish new colonies at considerable distances throughout the environment, therefore enabling extensive genetic mixing within the population (Thorne et al. 1999; Vargo et al. 2003; DeHeer and Vargo 2004). However, the presence of a single simple-family in our sampled colonies raises the question of whether the mixed families observed in our study are derived from colony foundation by the association of several primary reproductives, as observed in other higher termite species (Brandl et al. 2001; Thorne 1984). It also raises the question of how quickly primary reproductives are replaced by neotenics in the field (*i*.*e*., colonies becoming extended families), as well as the average colony age in the field, as neotenic reproduction occurs in older colonies, allowing them to extend their lifespan beyond those of the primary reproductives. This finding may also influence colony reproduction through dispersal, as long-lived organisms usually exhibit an older reproduction age. Therefore, both intra- and interspecific competition may also occur indirectly during colony foundation. Determining whether the timing of the nuptial flight of two species (*i*.*e*., time of the season and the colony age) and/or the ability to disperse (*i*.*e*., sex-ratio, weight and number of alates produced) and locate favorable nesting and foraging sites may provide further insights into the competition between these two termite species (Jones and Trosset 1991; Vargo and Husseneder 2009). Similarly, expanding our knowledge of the colony structure and breeding system of additional species of higher termites will definitively enhance our understanding of the mechanisms underlying the evolution of mating systems in termites.

## Conclusion

Interspecific competition is often hard to assess due to the presence of many indirect pressures in the environment, such as climate change, resource abundance, human activity, as well as shifts in niche roles (Kaplan and Denno 2007). Our study provides a first look at the breeding system of the higher termite species *A. parvulus* and offers insight as to how this termite species faces intraspecific competition and avoids interspecific competition with the lower termite *R. flavipes*. Colonies of the two species exhibit drastic differences in their foraging strategies, both in terms of habitat use (opened areas *vs*. forests), as well as in the size of their foraging range and the size of their foraging buffer with adjacent neighboring conspecific colonies. These differences are probably associated with differences in food preference, with *R. flavipes* feeding on large pieces of wood while *A. parvulus* feed on grass stems and small twigs, reducing their niche overlap and facilitating their co-existence across a shared urban landscape.

## Acknowledgments

We are grateful to Kyle Gilder for his help in the field. We thank BASF and the Urban Endowment Fund at Texas A& M University for providing funding for this work. This preprint was typeset with the bioRxiv word template by @Chrelli: www.github.com/chrelli/bioRxiv-word-template

## Data availability

The data presented in this study are available in Supplementary Materials and will be deposited on the Open Science Framework data depository (*https://osf.io/*) upon acceptance..

