## Supplemental Figure S1 for "Comparative genetic study of the colony structure and colony spatial distribution between the higher termite *Amitermes parvulus* and the lower, subterranean termite *Reticulitermes flavipes* in an urban environment"

| | PF1 | PF2 | PF3 | PF4 | PF5 | PF7 | PF8 | PF11 | PF67 | PF14 | PF15 | PF16 | PF17 | PF23 | PF26 | PF28 | PF50 | PF51 | PF96 | PF52 | PF53 | PF55 | PF56 | PF57 | PF66 | PF29 | PF34 | PF73 | PF74 | PF75 | PF77 | PF81 | PF82 | PF97 |
|---|---|---|---|---|---|---|---|---|---|---|---|---|---|---|---|---|---|---|---|---|---|---|---|---|---|---|---|---|---|---|---|---|---|---|
| 1 | 0 | 0.306 | 0.239 | 0.197 | 0.321 | 0.307 | 0.277 | 0.295 | 0.299 | 0.352 | 0.282 | 0.336 | 0.39 | 0.318 | 0.263 | 0.356 | 0.406 | 0.365 | 0.324 | 0.381 | 0.315 | 0.34 | 0.366 | 0.319 | 0.338 | 0.251 | 0.265 | 0.274 | 0.272 | 0.367 | 0.3 | 0.364 | 0.266 | 0.318 |
| 2 | 0.306 | 0 | 0.037 | 0.195 | 0.311 | 0.291 | 0.206 | 0.202 | 0.23 | 0.28 | 0.238 | 0.3 | 0.387 | 0.334 | 0.315 | 0.311 | 0.374 | 0.323 | 0.269 | 0.331 | 0.292 | 0.317 | 0.17 | 0.072 | 0.349 | 0.301 | 0.296 | 0.277 | 0.279 | 0.381 | 0.271 | 0.393 | 0.289 | 0.28 |
| 3 | 0.239 | 0.037 | 0 | 0.188 | 0.307 | 0.292 | 0.179 | 0.188 | 0.188 | 0.189 | 0.231 | 0.244 | 0.359 | 0.313 | 0.295 | 0.304 | 0.356 | 0.308 | 0.259 | 0.259 | 0.271 | 0.296 | 0.244 | 0.145 | 0.328 | 0.279 | 0.264 | 0.253 | 0.247 | 0.346 | 0.253 | 0.369 | 0.215 | 0.285 |
| 4 | 0.197 | 0.195 | 0.188 | 0 | 0.172 | 0.158 | 0.123 | 0.149 | 0.148 | 0.221 | 0.153 | 0.215 | 0.235 | 0.228 | 0.222 | 0.144 | 0.292 | 0.258 | 0.208 | 0.279 | 0.167 | 0.243 | 0.108 | 0.115 | 0.224 | 0.197 | 0.216 | 0.202 | 0.206 | 0.235 | 0.172 | 0.258 | 0.213 | 0.161 |
| 5 | 0.321 | 0.311 | 0.307 | 0.172 | 0 | -0.038 | 0.27 | 0.276 | 0.299 | 0.332 | 0.284 | 0.315 | 0.399 | 0.213 | 0.318 | 0.284 | 0.386 | 0.346 | 0.299 | 0.319 | 0.252 | 0.319 | 0.208 | 0.272 | 0.272 | 0.245 | 0.361 | 0.326 | 0.319 | 0.297 | 0.255 | 0.287 | 0.326 | 0.204 |
| 7 | 0.307 | 0.291 | 0.292 | 0.158 | -0.038 | 0 | 0.254 | 0.257 | 0.284 | 0.32 | 0.266 | 0.303 | 0.386 | 0.206 | 0.306 | 0.272 | 0.374 | 0.333 | 0.287 | 0.305 | 0.244 | 0.307 | 0.194 | 0.248 | 0.268 | 0.231 | 0.343 | 0.309 | 0.304 | 0.296 | 0.236 | 0.287 | 0.313 | 0.195 |
| 8 | 0.277 | 0.206 | 0.179 | 0.123 | 0.27 | 0.254 | 0 | -0.01 | -0.005 | 0.222 | 0.133 | 0.193 | 0.258 | 0.28 | 0.173 | 0.106 | 0.26 | 0.252 | 0.197 | 0.289 | 0.181 | 0.253 | 0.236 | 0.159 | 0.232 | 0.208 | 0.27 | 0.244 | 0.232 | 0.343 | 0.173 | 0.341 | 0.268 | 0.227 |
| 11 | 0.295 | 0.202 | 0.188 | 0.149 | 0.276 | 0.257 | -0.01 | 0 | 0.005 | 0.238 | 0.16 | 0.218 | 0.302 | 0.292 | 0.203 | 0.142 | 0.279 | 0.269 | 0.221 | 0.298 | 0.208 | 0.264 | 0.239 | 0.189 | 0.241 | 0.227 | 0.264 | 0.242 | 0.235 | 0.356 | 0.192 | 0.353 | 0.281 | 0.224 |
| 67 | 0.299 | 0.23 | 0.188 | 0.148 | 0.299 | 0.284 | -0.005 | 0.005 | 0 | 0.213 | 0.209 | 0.228 | 0.25 | 0.301 | 0.224 | 0.137 | 0.293 | 0.279 | 0.23 | 0.302 | 0.216 | 0.274 | 0.284 | 0.219 | 0.259 | 0.24 | 0.304 | 0.275 | 0.26 | 0.364 | 0.206 | 0.361 | 0.289 | 0.261 |
| 14 | 0.352 | 0.28 | 0.189 | 0.221 | 0.332 | 0.32 | 0.222 | 0.238 | 0.213 | 0 | 0.29 | 0.167 | 0.332 | 0.306 | 0.309 | 0.297 | 0.376 | 0.323 | 0.281 | 0.245 | 0.286 | 0.311 | 0.336 | 0.257 | 0.343 | 0.291 | 0.37 | 0.333 | 0.319 | 0.344 | 0.272 | 0.38 | 0.263 | 0.291 |
| 15 | 0.282 | 0.238 | 0.231 | 0.153 | 0.284 | 0.266 | 0.133 | 0.16 | 0.209 | 0.29 | 0 | 0.251 | 0.346 | 0.265 | 0.213 | 0.244 | 0.329 | 0.297 | 0.244 | 0.323 | 0.219 | 0.283 | 0.238 | 0.172 | 0.307 | 0.259 | 0.239 | 0.23 | 0.238 | 0.335 | 0.219 | 0.361 | 0.247 | 0.257 |
| 16 | 0.336 | 0.3 | 0.244 | 0.215 | 0.315 | 0.303 | 0.193 | 0.218 | 0.228 | 0.167 | 0.251 | 0 | 0.372 | 0.311 | 0.2 | 0.189 | 0.268 | 0.255 | 0.196 | 0.215 | 0.197 | 0.281 | 0.304 | 0.186 | 0.241 | 0.23 | 0.312 | 0.285 | 0.271 | 0.374 | 0.195 | 0.366 | 0.263 | 0.275 |
| 17 | 0.39 | 0.387 | 0.359 | 0.235 | 0.399 | 0.386 | 0.258 | 0.302 | 0.25 | 0.332 | 0.346 | 0.372 | 0 | 0.395 | 0.375 | 0.3 | 0.44 | 0.395 | 0.353 | 0.418 | 0.351 | 0.376 | 0.403 | 0.353 | 0.357 | 0.316 | 0.392 | 0.368 | 0.361 | 0.437 | 0.338 | 0.453 | 0.354 | 0.383 |
| 23 | 0.318 | 0.334 | 0.313 | 0.228 | 0.213 | 0.206 | 0.28 | 0.292 | 0.301 | 0.306 | 0.265 | 0.311 | 0.395 | 0 | 0.263 | 0.302 | 0.383 | 0.342 | 0.299 | 0.357 | 0.243 | 0.315 | 0.275 | 0.295 | 0.255 | 0.134 | 0.377 | 0.339 | 0.324 | 0.279 | 0.263 | 0.269 | 0.321 | 0.23 |
| 26 | 0.263 | 0.315 | 0.295 | 0.222 | 0.318 | 0.306 | 0.173 | 0.203 | 0.224 | 0.309 | 0.213 | 0.2 | 0.375 | 0.263 | 0 | 0.184 | 0.287 | 0.282 | 0.227 | 0.273 | 0.212 | 0.259 | 0.309 | 0.219 | 0.202 | 0.132 | 0.321 | 0.293 | 0.279 | 0.376 | 0.207 | 0.302 | 0.302 | 0.242 |
| 28 | 0.356 | 0.311 | 0.304 | 0.144 | 0.284 | 0.272 | 0.106 | 0.142 | 0.137 | 0.297 | 0.244 | 0.189 | 0.3 | 0.302 | 0.184 | 0 | 0.248 | 0.272 | 0.208 | 0.346 | 0.158 | 0.295 | 0.235 | 0.202 | 0.166 | 0.227 | 0.296 | 0.277 | 0.268 | 0.339 | 0.176 | 0.333 | 0.319 | 0.208 |
| 50 | 0.406 | 0.374 | 0.356 | 0.292 | 0.386 | 0.374 | 0.26 | 0.279 | 0.293 | 0.376 | 0.329 | 0.268 | 0.44 | 0.383 | 0.287 | 0.248 | 0 | 0.009 | 0.009 | 0.374 | 0.25 | 0.348 | 0.369 | 0.29 | 0.263 | 0.272 | 0.37 | 0.347 | 0.332 | 0.438 | 0.26 | 0.409 | 0.244 | 0.338 |
| 51 | 0.365 | 0.323 | 0.308 | 0.258 | 0.346 | 0.333 | 0.252 | 0.269 | 0.279 | 0.323 | 0.297 | 0.255 | 0.395 | 0.342 | 0.282 | 0.272 | 0.009 | 0 | -0.029 | 0.336 | 0.223 | 0.315 | 0.32 | 0.251 | 0.279 | 0.256 | 0.357 | 0.326 | 0.312 | 0.388 | 0.25 | 0.333 | 0.204 | 0.313 |
| 96 | 0.324 | 0.269 | 0.259 | 0.208 | 0.299 | 0.287 | 0.197 | 0.221 | 0.23 | 0.281 | 0.244 | 0.196 | 0.353 | 0.299 | 0.227 | 0.208 | 0.009 | -0.029 | 0 | 0.293 | 0.169 | 0.273 | 0.267 | 0.165 | 0.228 | 0.194 | 0.313 | 0.283 | 0.269 | 0.348 | 0.197 | 0.3 | 0.178 | 0.271 |
| 52 | 0.381 | 0.331 | 0.259 | 0.279 | 0.319 | 0.305 | 0.289 | 0.298 | 0.302 | 0.245 | 0.323 | 0.215 | 0.418 | 0.357 | 0.273 | 0.346 | 0.374 | 0.336 | 0.293 | 0 | 0.31 | 0.337 | 0.365 | 0.256 | 0.346 | 0.305 | 0.395 | 0.359 | 0.345 | 0.419 | 0.296 | 0.41 | 0.269 | 0.343 |
| 53 | 0.315 | 0.292 | 0.271 | 0.167 | 0.252 | 0.244 | 0.181 | 0.208 | 0.216 | 0.286 | 0.219 | 0.197 | 0.351 | 0.243 | 0.212 | 0.158 | 0.25 | 0.223 | 0.169 | 0.31 | 0 | 0.18 | 0.217 | 0.19 | 0.191 | 0.203 | 0.303 | 0.273 | 0.259 | 0.281 | 0.172 | 0.225 | 0.207 | 0.205 |
| 55 | 0.34 | 0.317 | 0.296 | 0.243 | 0.319 | 0.307 | 0.253 | 0.264 | 0.274 | 0.311 | 0.283 | 0.281 | 0.376 | 0.315 | 0.259 | 0.295 | 0.348 | 0.315 | 0.273 | 0.337 | 0.18 | 0 | 0.324 | 0.268 | 0.31 | 0.272 | 0.325 | 0.266 | 0.246 | 0.378 | 0.207 | 0.368 | 0.228 | 0.297 |
| 56 | 0.366 | 0.17 | 0.244 | 0.108 | 0.208 | 0.194 | 0.236 | 0.239 | 0.284 | 0.336 | 0.238 | 0.304 | 0.403 | 0.275 | 0.309 | 0.235 | 0.369 | 0.32 | 0.267 | 0.365 | 0.217 | 0.324 | 0 | 0.104 | 0.258 | 0.28 | 0.335 | 0.304 | 0.307 | 0.268 | 0.243 | 0.239 | 0.33 | 0.135 |
| 57 | 0.319 | 0.072 | 0.145 | 0.115 | 0.272 | 0.248 | 0.159 | 0.189 | 0.219 | 0.257 | 0.172 | 0.186 | 0.353 | 0.295 | 0.219 | 0.202 | 0.29 | 0.251 | 0.165 | 0.256 | 0.19 | 0.268 | 0.104 | 0 | 0.281 | 0.212 | 0.309 | 0.268 | 0.262 | 0.376 | 0.181 | 0.363 | 0.252 | 0.243 |
| 66 | 0.338 | 0.349 | 0.328 | 0.224 | 0.272 | 0.268 | 0.232 | 0.241 | 0.259 | 0.343 | 0.307 | 0.241 | 0.357 | 0.255 | 0.202 | 0.166 | 0.263 | 0.279 | 0.228 | 0.346 | 0.191 | 0.31 | 0.258 | 0.281 | 0 | 0.053 | 0.325 | 0.305 | 0.29 | 0.309 | 0.21 | 0.254 | 0.312 | 0.166 |
| 29 | 0.251 | 0.301 | 0.279 | 0.197 | 0.245 | 0.231 | 0.208 | 0.227 | 0.24 | 0.291 | 0.259 | 0.23 | 0.316 | 0.134 | 0.132 | 0.227 | 0.272 | 0.256 | 0.194 | 0.305 | 0.203 | 0.272 | 0.28 | 0.212 | 0.053 | 0 | 0.321 | 0.285 | 0.265 | 0.361 | 0.193 | 0.267 | 0.26 | 0.204 |
| 34 | 0.265 | 0.296 | 0.264 | 0.216 | 0.361 | 0.343 | 0.27 | 0.264 | 0.304 | 0.37 | 0.239 | 0.312 | 0.392 | 0.377 | 0.321 | 0.296 | 0.37 | 0.357 | 0.313 | 0.395 | 0.303 | 0.325 | 0.335 | 0.309 | 0.325 | 0.321 | 0 | 0.011 | 0.051 | 0.354 | 0.283 | 0.435 | 0.238 | 0.31 |
| 73 | 0.274 | 0.277 | 0.253 | 0.202 | 0.326 | 0.309 | 0.244 | 0.242 | 0.275 | 0.333 | 0.23 | 0.285 | 0.368 | 0.339 | 0.293 | 0.277 | 0.347 | 0.326 | 0.283 | 0.359 | 0.273 | 0.266 | 0.304 | 0.268 | 0.305 | 0.285 | 0.011 | 0 | [illegible] | 0.344 | 0.254 | 0.399 | 0.243 | 0.283 |
| 74 | 0.272 | 0.279 | 0.247 | 0.206 | 0.319 | 0.304 | 0.232 | 0.235 | 0.26 | 0.319 | 0.238 | 0.271 | 0.361 | 0.324 | 0.279 | 0.268 | 0.332 | 0.312 | 0.269 | 0.345 | 0.259 | 0.246 | 0.307 | 0.262 | 0.29 | 0.265 | 0.051 | [illegible] | 0 | 0.341 | 0.242 | 0.385 | 0.245 | 0.28 |
| 75 | 0.367 | 0.381 | 0.346 | 0.235 | 0.297 | 0.296 | 0.343 | 0.356 | 0.364 | 0.344 | 0.335 | 0.374 | 0.437 | 0.279 | 0.376 | 0.339 | 0.438 | 0.388 | 0.348 | 0.419 | 0.281 | 0.378 | 0.268 | 0.376 | 0.309 | 0.361 | 0.354 | 0.344 | 0.341 | 0 | 0.317 | 0.263 | 0.334 | 0.251 |
| 77 | 0.3 | 0.271 | 0.253 | 0.172 | 0.255 | 0.236 | 0.173 | 0.192 | 0.206 | 0.272 | 0.219 | 0.195 | 0.338 | 0.263 | 0.207 | 0.176 | 0.26 | 0.25 | 0.197 | 0.296 | 0.172 | 0.207 | 0.243 | 0.181 | 0.21 | 0.193 | 0.283 | 0.254 | 0.242 | 0.317 | 0 | 0.314 | 0.263 | 0.218 |
| 81 | 0.364 | 0.393 | 0.369 | 0.258 | 0.287 | 0.287 | 0.341 | 0.353 | 0.361 | 0.38 | 0.361 | 0.366 | 0.453 | 0.269 | 0.302 | 0.333 | 0.409 | 0.333 | 0.3 | 0.41 | 0.225 | 0.368 | 0.239 | 0.363 | 0.254 | 0.267 | 0.435 | 0.399 | 0.385 | 0.263 | 0.314 | 0 | 0.366 | 0.198 |
| 82 | 0.266 | 0.289 | 0.215 | 0.213 | 0.326 | 0.313 | 0.268 | 0.281 | 0.289 | 0.263 | 0.247 | 0.263 | 0.354 | 0.321 | 0.302 | 0.319 | 0.244 | 0.204 | 0.178 | 0.269 | 0.207 | 0.228 | 0.33 | 0.252 | 0.312 | 0.26 | 0.238 | 0.243 | 0.245 | 0.334 | 0.263 | 0.366 | 0 | 0.31 |
| 97 | 0.318 | 0.28 | 0.285 | 0.161 | 0.204 | 0.195 | 0.227 | 0.224 | 0.261 | 0.291 | 0.257 | 0.275 | 0.383 | 0.23 | 0.242 | 0.208 | 0.338 | 0.313 | 0.271 | 0.343 | 0.205 | 0.297 | 0.135 | 0.243 | 0.166 | 0.204 | 0.31 | 0.283 | 0.28 | 0.251 | 0.218 | 0.198 | 0.31 | 0 |

**Figure S1**: $F_{ST}$ genetic differentiation values between each pair of stations are visualized using a station- by- station matrix for Reticulitermes flavipes samples. Cells are colored according to the genetic differentiation between each pair of stations; darker shades of grey indicate that stations are more genetically similar to each other (i.e., more likely to belong to the same colony).
