## Supplemental Figure S2 for "Comparative genetic study of the colony structure and colony spatial distribution between the higher termite *Amitermes parvulus* and the lower, subterranean termite *Reticulitermes flavipes* in an urban environment"

| | Large-scale experiment | | | | | | | | | | | | | | | | | | | | | |
|---|---|---|---|---|---|---|---|---|---|---|---|---|---|---|---|---|---|---|---|---|---|---|
| **Large-scale experiment** | | | | | | | | | | | | | | | | | | | | | | |
| *Am1* | 0.00 | 0.02 | 0.36 | 0.39 | 0.31 | 0.35 | 0.38 | 0.34 | 0.37 | 0.34 | 0.32 | 0.37 | 0.34 | 0.41 | 0.42 | 0.35 | 0.29 | 0.34 | 0.47 | 0.26 | 0.29 | 0.40 |
| *Am2* | 0.02 | 0.00 | 0.39 | 0.41 | 0.34 | 0.38 | 0.40 | 0.36 | 0.36 | 0.33 | 0.32 | 0.38 | 0.32 | 0.43 | 0.44 | 0.37 | 0.33 | 0.36 | 0.45 | 0.28 | 0.32 | 0.41 |
| *Am9* | 0.36 | 0.39 | 0.00 | 0.15 | 0.04 | 0.46 | 0.47 | 0.44 | 0.48 | 0.46 | 0.43 | 0.45 | 0.46 | 0.53 | 0.53 | 0.38 | 0.36 | 0.46 | 0.53 | 0.38 | 0.44 | 0.50 |
| *Am10* | 0.39 | 0.41 | 0.15 | 0.00 | 0.06 | 0.47 | 0.48 | 0.45 | 0.47 | 0.43 | 0.43 | 0.46 | 0.46 | 0.43 | 0.44 | 0.39 | 0.33 | 0.39 | 0.54 | 0.37 | 0.44 | 0.52 |
| *Am11* | 0.31 | 0.34 | 0.04 | 0.06 | 0.00 | 0.41 | 0.43 | 0.39 | 0.42 | 0.38 | 0.38 | 0.40 | 0.40 | 0.41 | 0.42 | 0.31 | 0.29 | 0.34 | 0.47 | 0.31 | 0.40 | 0.45 |
| *Am4* | 0.35 | 0.38 | 0.46 | 0.47 | 0.41 | 0.00 | 0.08 | 0.00 | 0.01 | 0.28 | 0.25 | 0.35 | 0.27 | 0.48 | 0.48 | 0.19 | 0.38 | 0.35 | 0.55 | 0.31 | 0.43 | 0.41 |
| *Am5* | 0.38 | 0.40 | 0.47 | 0.48 | 0.43 | 0.08 | 0.00 | 0.02 | 0.04 | 0.36 | 0.34 | 0.40 | 0.38 | 0.53 | 0.53 | 0.21 | 0.40 | 0.39 | 0.55 | 0.34 | 0.45 | 0.45 |
| *Am12* | 0.34 | 0.36 | 0.44 | 0.45 | 0.39 | 0.00 | 0.02 | 0.00 | -0.01 | 0.31 | 0.30 | 0.34 | 0.30 | 0.49 | 0.50 | 0.19 | 0.38 | 0.35 | 0.53 | 0.31 | 0.42 | 0.45 |
| *Am13* | 0.37 | 0.36 | 0.48 | 0.47 | 0.42 | 0.01 | 0.04 | -0.01 | 0.00 | 0.36 | 0.33 | 0.36 | 0.35 | 0.53 | 0.53 | 0.18 | 0.38 | 0.38 | 0.55 | 0.31 | 0.43 | 0.44 |
| *Am14* | 0.34 | 0.33 | 0.46 | 0.43 | 0.38 | 0.28 | 0.36 | 0.31 | 0.36 | 0.00 | 0.00 | 0.04 | 0.02 | 0.44 | 0.47 | 0.27 | 0.39 | 0.35 | 0.55 | 0.26 | 0.46 | 0.42 |
| *Am15* | 0.32 | 0.32 | 0.43 | 0.43 | 0.38 | 0.25 | 0.34 | 0.30 | 0.33 | 0.00 | 0.00 | 0.05 | 0.03 | 0.46 | 0.47 | 0.24 | 0.40 | 0.37 | 0.55 | 0.25 | 0.45 | 0.40 |
| *Am7* | 0.37 | 0.38 | 0.45 | 0.46 | 0.40 | 0.35 | 0.40 | 0.34 | 0.36 | 0.04 | 0.05 | 0.00 | 0.00 | 0.51 | 0.54 | 0.30 | 0.42 | 0.43 | 0.58 | 0.34 | 0.49 | 0.50 |
| *Am8* | 0.34 | 0.32 | 0.46 | 0.46 | 0.40 | 0.27 | 0.38 | 0.30 | 0.35 | 0.02 | 0.03 | 0.00 | 0.00 | 0.50 | 0.52 | 0.30 | 0.42 | 0.43 | 0.55 | 0.29 | 0.47 | 0.43 |
| *Am16* | 0.41 | 0.43 | 0.53 | 0.43 | 0.41 | 0.48 | 0.53 | 0.49 | 0.53 | 0.44 | 0.46 | 0.51 | 0.50 | 0.00 | 0.00 | 0.57 | 0.39 | 0.40 | 0.64 | 0.39 | 0.47 | 0.51 |
| *Am17* | 0.42 | 0.44 | 0.53 | 0.44 | 0.42 | 0.48 | 0.53 | 0.50 | 0.53 | 0.47 | 0.47 | 0.54 | 0.52 | 0.00 | 0.00 | 0.53 | 0.41 | 0.38 | 0.63 | 0.40 | 0.48 | 0.50 |
| *Am18* | 0.35 | 0.37 | 0.38 | 0.39 | 0.31 | 0.19 | 0.21 | 0.19 | 0.18 | 0.27 | 0.24 | 0.30 | 0.30 | 0.57 | 0.53 | 0.00 | 0.31 | 0.34 | 0.54 | 0.24 | 0.42 | 0.42 |
| *Am19* | 0.29 | 0.33 | 0.36 | 0.33 | 0.29 | 0.38 | 0.40 | 0.38 | 0.38 | 0.39 | 0.40 | 0.42 | 0.42 | 0.39 | 0.41 | 0.31 | 0.00 | 0.26 | 0.53 | 0.26 | 0.33 | 0.42 |
| *Am20* | 0.34 | 0.36 | 0.46 | 0.39 | 0.34 | 0.35 | 0.39 | 0.35 | 0.38 | 0.35 | 0.37 | 0.43 | 0.43 | 0.40 | 0.38 | 0.34 | 0.26 | 0.00 | 0.47 | 0.24 | 0.38 | 0.36 |
| *Am21* | 0.47 | 0.45 | 0.53 | 0.54 | 0.47 | 0.55 | 0.55 | 0.53 | 0.55 | 0.55 | 0.55 | 0.58 | 0.55 | 0.64 | 0.63 | 0.54 | 0.53 | 0.47 | 0.00 | 0.49 | 0.56 | 0.58 |
| *Am22* | 0.26 | 0.28 | 0.38 | 0.37 | 0.31 | 0.31 | 0.34 | 0.31 | 0.31 | 0.26 | 0.25 | 0.34 | 0.29 | 0.39 | 0.40 | 0.24 | 0.26 | 0.24 | 0.49 | 0.00 | 0.37 | 0.39 |
| *Am23* | 0.29 | 0.32 | 0.44 | 0.44 | 0.40 | 0.43 | 0.45 | 0.42 | 0.43 | 0.46 | 0.45 | 0.49 | 0.47 | 0.47 | 0.48 | 0.42 | 0.33 | 0.38 | 0.56 | 0.37 | 0.00 | 0.44 |
| *Am25* | 0.40 | 0.41 | 0.50 | 0.52 | 0.45 | 0.41 | 0.45 | 0.45 | 0.44 | 0.42 | 0.40 | 0.50 | 0.43 | 0.51 | 0.50 | 0.42 | 0.42 | 0.36 | 0.58 | 0.39 | 0.44 | 0.00 |
| **Fine-scale experiment** | | | | | | | | | | | | | | | | | | | | | | |
| *PF54* | 0.51 | 0.51 | 0.55 | 0.54 | 0.52 | 0.60 | 0.64 | 0.62 | 0.63 | 0.62 | 0.63 | 0.65 | 0.64 | 0.60 | 0.59 | 0.58 | 0.57 | 0.56 | 0.63 | 0.57 | 0.59 | 0.58 |
| *PF41* | 0.40 | 0.44 | 0.48 | 0.47 | 0.43 | 0.47 | 0.49 | 0.47 | 0.47 | 0.47 | 0.47 | 0.51 | 0.49 | 0.51 | 0.51 | 0.44 | 0.41 | 0.34 | 0.45 | 0.39 | 0.45 | 0.32 |
| *PF58* | 0.43 | 0.42 | 0.58 | 0.58 | 0.51 | 0.48 | 0.50 | 0.51 | 0.52 | 0.46 | 0.45 | 0.51 | 0.46 | 0.62 | 0.62 | 0.48 | 0.49 | 0.39 | 0.57 | 0.39 | 0.52 | 0.28 |
| *PF59* | 0.38 | 0.41 | 0.46 | 0.46 | 0.41 | 0.43 | 0.45 | 0.44 | 0.44 | 0.43 | 0.42 | 0.48 | 0.45 | 0.49 | 0.49 | 0.42 | 0.37 | 0.30 | 0.46 | 0.34 | 0.45 | 0.26 |
| *PF61* | 0.42 | 0.45 | 0.50 | 0.49 | 0.44 | 0.47 | 0.49 | 0.49 | 0.50 | 0.47 | 0.48 | 0.51 | 0.49 | 0.53 | 0.53 | 0.46 | 0.42 | 0.35 | 0.49 | 0.39 | 0.47 | 0.27 |
| *PF63* | 0.39 | 0.40 | 0.47 | 0.47 | 0.41 | 0.44 | 0.47 | 0.46 | 0.46 | 0.46 | 0.46 | 0.50 | 0.48 | 0.51 | 0.51 | 0.41 | 0.39 | 0.33 | 0.45 | 0.38 | 0.44 | 0.22 |
| *PF65* | 0.42 | 0.39 | 0.57 | 0.57 | 0.51 | 0.49 | 0.53 | 0.51 | 0.52 | 0.50 | 0.49 | 0.53 | 0.50 | 0.58 | 0.58 | 0.48 | 0.42 | 0.43 | 0.52 | 0.40 | 0.47 | 0.31 |
| *PF68* | 0.38 | 0.39 | 0.44 | 0.46 | 0.40 | 0.40 | 0.43 | 0.42 | 0.41 | 0.43 | 0.43 | 0.47 | 0.45 | 0.50 | 0.50 | 0.37 | 0.38 | 0.31 | 0.46 | 0.35 | 0.42 | 0.22 |
| *PF69* | 0.40 | 0.39 | 0.49 | 0.49 | 0.44 | 0.46 | 0.49 | 0.48 | 0.48 | 0.46 | 0.47 | 0.49 | 0.47 | 0.54 | 0.53 | 0.45 | 0.42 | 0.37 | 0.42 | 0.39 | 0.46 | 0.28 |
| *PF70* | 0.37 | 0.38 | 0.42 | 0.42 | 0.36 | 0.44 | 0.46 | 0.44 | 0.44 | 0.42 | 0.43 | 0.46 | 0.44 | 0.49 | 0.48 | 0.40 | 0.39 | 0.32 | 0.42 | 0.35 | 0.44 | 0.28 |
| *PF76* | 0.41 | 0.43 | 0.49 | 0.50 | 0.44 | 0.44 | 0.48 | 0.47 | 0.46 | 0.45 | 0.45 | 0.50 | 0.46 | 0.52 | 0.52 | 0.42 | 0.42 | 0.34 | 0.48 | 0.38 | 0.46 | 0.23 |
| *PF79* | 0.41 | 0.42 | 0.51 | 0.51 | 0.45 | 0.47 | 0.50 | 0.49 | 0.49 | 0.48 | 0.49 | 0.52 | 0.50 | 0.56 | 0.55 | 0.44 | 0.38 | 0.38 | 0.45 | 0.42 | 0.47 | 0.26 |
| *PF84* | 0.39 | 0.41 | 0.48 | 0.48 | 0.43 | 0.47 | 0.49 | 0.49 | 0.49 | 0.45 | 0.45 | 0.49 | 0.47 | 0.52 | 0.52 | 0.46 | 0.43 | 0.35 | 0.47 | 0.38 | 0.47 | 0.27 |
| *PF87* | 0.38 | 0.38 | 0.46 | 0.46 | 0.41 | 0.43 | 0.45 | 0.45 | 0.45 | 0.41 | 0.41 | 0.45 | 0.42 | 0.49 | 0.49 | 0.41 | 0.38 | 0.31 | 0.42 | 0.35 | 0.43 | 0.25 |
| *PF89* | 0.36 | 0.38 | 0.45 | 0.45 | 0.40 | 0.43 | 0.46 | 0.45 | 0.46 | 0.42 | 0.42 | 0.46 | 0.44 | 0.50 | 0.50 | 0.44 | 0.39 | 0.34 | 0.44 | 0.36 | 0.43 | 0.22 |
| *PF90* | 0.36 | 0.37 | 0.42 | 0.44 | 0.38 | 0.42 | 0.44 | 0.43 | 0.42 | 0.42 | 0.41 | 0.46 | 0.44 | 0.48 | 0.48 | 0.37 | 0.37 | 0.30 | 0.41 | 0.32 | 0.42 | 0.24 |
| *PF91* | 0.38 | 0.38 | 0.44 | 0.45 | 0.40 | 0.42 | 0.45 | 0.44 | 0.44 | 0.42 | 0.42 | 0.46 | 0.43 | 0.49 | 0.49 | 0.40 | 0.37 | 0.33 | 0.39 | 0.36 | 0.43 | 0.24 |
| *PF92* | 0.37 | 0.38 | 0.43 | 0.44 | 0.39 | 0.42 | 0.45 | 0.43 | 0.43 | 0.40 | 0.40 | 0.45 | 0.41 | 0.48 | 0.48 | 0.38 | 0.38 | 0.29 | 0.44 | 0.32 | 0.43 | 0.23 |
| *PF93* | 0.35 | 0.37 | 0.45 | 0.44 | 0.39 | 0.38 | 0.42 | 0.41 | 0.41 | 0.40 | 0.41 | 0.45 | 0.42 | 0.47 | 0.47 | 0.35 | 0.36 | 0.28 | 0.43 | 0.33 | 0.42 | 0.20 |
| *PF94* | 0.39 | 0.40 | 0.44 | 0.45 | 0.40 | 0.42 | 0.45 | 0.44 | 0.44 | 0.43 | 0.43 | 0.48 | 0.45 | 0.49 | 0.49 | 0.39 | 0.38 | 0.30 | 0.43 | 0.34 | 0.43 | 0.19 |
| *PF95* | 0.36 | 0.36 | 0.44 | 0.43 | 0.38 | 0.40 | 0.43 | 0.42 | 0.43 | 0.41 | 0.41 | 0.45 | 0.42 | 0.47 | 0.47 | 0.35 | 0.36 | 0.28 | 0.39 | 0.34 | 0.41 | 0.24 |

| | Fine-scale experiment | | | | | | | | | | | | | | | | | | | | |
|---|---|---|---|---|---|---|---|---|---|---|---|---|---|---|---|---|---|---|---|---|---|
| **Large-scale experiment** | | | | | | | | | | | | | | | | | | | | | |
| *Am1* | 0.51 | 0.40 | 0.43 | 0.38 | 0.42 | 0.39 | 0.42 | 0.38 | 0.40 | 0.37 | 0.41 | 0.41 | 0.39 | 0.38 | 0.36 | 0.36 | 0.38 | 0.37 | 0.35 | 0.39 | 0.36 |
| *Am2* | 0.51 | 0.44 | 0.42 | 0.41 | 0.45 | 0.40 | 0.39 | 0.39 | 0.39 | 0.38 | 0.43 | 0.42 | 0.41 | 0.38 | 0.38 | 0.37 | 0.38 | 0.38 | 0.37 | 0.40 | 0.36 |
| *Am9* | 0.55 | 0.48 | 0.58 | 0.46 | 0.50 | 0.47 | 0.57 | 0.44 | 0.49 | 0.42 | 0.49 | 0.51 | 0.48 | 0.46 | 0.45 | 0.42 | 0.44 | 0.43 | 0.45 | 0.44 | 0.44 |
| *Am10* | 0.54 | 0.47 | 0.58 | 0.46 | 0.49 | 0.47 | 0.57 | 0.46 | 0.49 | 0.42 | 0.50 | 0.51 | 0.48 | 0.46 | 0.45 | 0.44 | 0.45 | 0.44 | 0.44 | 0.45 | 0.43 |
| *Am11* | 0.52 | 0.43 | 0.51 | 0.41 | 0.44 | 0.41 | 0.51 | 0.40 | 0.44 | 0.36 | 0.44 | 0.45 | 0.43 | 0.41 | 0.40 | 0.38 | 0.40 | 0.39 | 0.39 | 0.40 | 0.38 |
| *Am4* | 0.60 | 0.47 | 0.48 | 0.43 | 0.47 | 0.44 | 0.49 | 0.40 | 0.46 | 0.44 | 0.44 | 0.47 | 0.47 | 0.43 | 0.43 | 0.42 | 0.42 | 0.42 | 0.38 | 0.42 | 0.40 |
| *Am5* | 0.64 | 0.49 | 0.50 | 0.45 | 0.49 | 0.47 | 0.53 | 0.43 | 0.49 | 0.46 | 0.48 | 0.50 | 0.49 | 0.45 | 0.46 | 0.44 | 0.45 | 0.45 | 0.42 | 0.45 | 0.43 |
| *Am12* | 0.62 | 0.47 | 0.51 | 0.44 | 0.49 | 0.46 | 0.51 | 0.42 | 0.48 | 0.44 | 0.47 | 0.49 | 0.49 | 0.45 | 0.45 | 0.43 | 0.44 | 0.43 | 0.41 | 0.44 | 0.42 |
| *Am13* | 0.63 | 0.47 | 0.52 | 0.44 | 0.50 | 0.46 | 0.52 | 0.41 | 0.48 | 0.44 | 0.46 | 0.49 | 0.49 | 0.45 | 0.46 | 0.42 | 0.44 | 0.43 | 0.41 | 0.44 | 0.43 |
| *Am14* | 0.62 | 0.47 | 0.46 | 0.43 | 0.47 | 0.46 | 0.50 | 0.43 | 0.46 | 0.42 | 0.45 | 0.48 | 0.45 | 0.41 | 0.42 | 0.42 | 0.42 | 0.40 | 0.40 | 0.43 | 0.41 |
| *Am15* | 0.63 | 0.47 | 0.45 | 0.42 | 0.48 | 0.46 | 0.49 | 0.43 | 0.47 | 0.43 | 0.45 | 0.49 | 0.45 | 0.41 | 0.42 | 0.41 | 0.42 | 0.40 | 0.41 | 0.43 | 0.41 |
| *Am7* | 0.65 | 0.51 | 0.51 | 0.48 | 0.51 | 0.50 | 0.53 | 0.47 | 0.49 | 0.46 | 0.50 | 0.52 | 0.49 | 0.45 | 0.46 | 0.46 | 0.46 | 0.45 | 0.45 | 0.48 | 0.45 |
| *Am8* | 0.64 | 0.49 | 0.46 | 0.45 | 0.49 | 0.48 | 0.50 | 0.45 | 0.47 | 0.44 | 0.46 | 0.50 | 0.47 | 0.42 | 0.44 | 0.44 | 0.43 | 0.41 | 0.42 | 0.45 | 0.42 |
| *Am16* | 0.60 | 0.51 | 0.62 | 0.49 | 0.53 | 0.51 | 0.58 | 0.50 | 0.54 | 0.49 | 0.52 | 0.56 | 0.52 | 0.49 | 0.50 | 0.48 | 0.49 | 0.48 | 0.47 | 0.49 | 0.47 |
| *Am17* | 0.59 | 0.51 | 0.62 | 0.49 | 0.53 | 0.51 | 0.58 | 0.50 | 0.53 | 0.48 | 0.52 | 0.55 | 0.52 | 0.49 | 0.50 | 0.48 | 0.49 | 0.48 | 0.47 | 0.49 | 0.47 |
| *Am18* | 0.58 | 0.44 | 0.48 | 0.42 | 0.46 | 0.41 | 0.48 | 0.37 | 0.45 | 0.40 | 0.42 | 0.44 | 0.46 | 0.41 | 0.44 | 0.37 | 0.40 | 0.38 | 0.35 | 0.39 | 0.35 |
| *Am19* | 0.57 | 0.41 | 0.49 | 0.37 | 0.42 | 0.39 | 0.42 | 0.38 | 0.42 | 0.39 | 0.42 | 0.38 | 0.43 | 0.38 | 0.39 | 0.37 | 0.37 | 0.38 | 0.36 | 0.38 | 0.36 |
| *Am20* | 0.56 | 0.34 | 0.39 | 0.30 | 0.35 | 0.33 | 0.43 | 0.31 | 0.37 | 0.32 | 0.34 | 0.38 | 0.35 | 0.31 | 0.34 | 0.30 | 0.33 | 0.29 | 0.28 | 0.30 | 0.28 |
| *Am21* | 0.63 | 0.45 | 0.57 | 0.46 | 0.49 | 0.45 | 0.52 | 0.46 | 0.42 | 0.42 | 0.48 | 0.45 | 0.47 | 0.42 | 0.44 | 0.41 | 0.39 | 0.44 | 0.43 | 0.43 | 0.39 |
| *Am22* | 0.57 | 0.39 | 0.39 | 0.34 | 0.39 | 0.38 | 0.40 | 0.35 | 0.39 | 0.35 | 0.38 | 0.42 | 0.38 | 0.35 | 0.36 | 0.32 | 0.36 | 0.32 | 0.33 | 0.34 | 0.34 |
| *Am23* | 0.59 | 0.45 | 0.52 | 0.45 | 0.47 | 0.44 | 0.47 | 0.42 | 0.46 | 0.44 | 0.46 | 0.47 | 0.47 | 0.43 | 0.43 | 0.42 | 0.43 | 0.43 | 0.42 | 0.43 | 0.41 |
| *Am25* | 0.58 | 0.32 | 0.28 | 0.26 | 0.27 | 0.22 | 0.31 | 0.22 | 0.28 | 0.28 | 0.23 | 0.26 | 0.27 | 0.25 | 0.22 | 0.24 | 0.24 | 0.23 | 0.20 | 0.19 | 0.24 |
| **Fine-scale experiment** | | | | | | | | | | | | | | | | | | | | | |
| *PF54* | 0.00 | 0.59 | 0.75 | 0.59 | 0.65 | 0.58 | 0.64 | 0.60 | 0.63 | 0.60 | 0.60 | 0.64 | 0.58 | 0.57 | 0.60 | 0.56 | 0.58 | 0.57 | 0.55 | 0.57 | 0.55 |
| *PF41* | 0.59 | 0.00 | 0.27 | 0.00 | 0.06 | 0.00 | 0.19 | 0.11 | 0.08 | 0.07 | 0.03 | 0.11 | 0.05 | 0.05 | 0.03 | 0.01 | 0.04 | 0.04 | 0.07 | 0.03 | 0.04 |
| *PF58* | 0.75 | 0.27 | 0.00 | 0.22 | 0.23 | 0.20 | 0.06 | 0.22 | 0.13 | 0.18 | 0.19 | 0.08 | 0.13 | 0.09 | 0.10 | 0.15 | 0.13 | 0.16 | 0.11 | 0.18 | 0.12 |
| *PF59* | 0.59 | 0.00 | 0.22 | 0.00 | -0.02 | 0.00 | 0.16 | 0.08 | 0.05 | 0.05 | 0.00 | 0.07 | 0.01 | 0.00 | 0.02 | -0.02 | -0.01 | -0.04 | 0.01 | -0.03 | 0.03 |
| *PF61* | 0.65 | 0.06 | 0.23 | -0.02 | 0.00 | 0.05 | 0.13 | 0.14 | 0.05 | 0.05 | 0.01 | 0.09 | 0.02 | 0.02 | 0.04 | 0.03 | 0.05 | 0.00 | 0.03 | -0.01 | 0.04 |
| *PF63* | 0.58 | 0.00 | 0.20 | 0.00 | 0.05 | 0.00 | 0.12 | 0.02 | 0.02 | 0.05 | 0.00 | 0.06 | 0.04 | 0.00 | 0.03 | -0.03 | 0.00 | 0.01 | -0.01 | -0.01 | 0.00 |
| *PF65* | 0.64 | 0.19 | 0.06 | 0.16 | 0.13 | 0.12 | 0.00 | 0.21 | 0.00 | 0.10 | 0.10 | -0.01 | 0.09 | 0.03 | 0.07 | 0.08 | 0.09 | 0.11 | 0.07 | 0.12 | 0.09 |
| *PF68* | 0.60 | 0.11 | 0.22 | 0.08 | 0.14 | 0.02 | 0.21 | 0.00 | 0.13 | 0.10 | 0.06 | 0.17 | 0.16 | 0.08 | 0.12 | 0.03 | 0.07 | 0.10 | 0.03 | 0.02 | 0.06 |
| *PF69* | 0.63 | 0.08 | 0.13 | 0.05 | 0.05 | 0.02 | 0.00 | 0.13 | 0.00 | 0.04 | 0.02 | 0.00 | 0.05 | -0.01 | 0.03 | 0.01 | 0.01 | 0.06 | 0.01 | 0.04 | 0.00 |
| *PF70* | 0.60 | 0.07 | 0.18 | 0.05 | 0.05 | 0.05 | 0.10 | 0.10 | 0.04 | 0.00 | 0.07 | 0.05 | 0.03 | 0.01 | 0.00 | 0.00 | 0.07 | 0.03 | 0.03 | 0.04 | 0.03 |
| *PF76* | 0.60 | 0.03 | 0.19 | 0.00 | 0.01 | 0.00 | 0.10 | 0.06 | 0.02 | 0.07 | 0.00 | 0.00 | 0.05 | 0.01 | 0.05 | 0.00 | 0.03 | 0.01 | -0.01 | -0.03 | 0.02 |
| *PF79* | 0.64 | 0.11 | 0.08 | 0.07 | 0.09 | 0.06 | -0.01 | 0.17 | 0.00 | 0.05 | 0.08 | 0.00 | 0.09 | 0.04 | 0.02 | 0.05 | 0.03 | 0.11 | 0.05 | 0.08 | 0.07 |
| *PF84* | 0.58 | 0.05 | 0.13 | 0.01 | 0.02 | 0.04 | 0.09 | 0.16 | 0.05 | 0.03 | 0.05 | 0.09 | 0.00 | -0.01 | -0.03 | 0.00 | 0.04 | -0.02 | 0.04 | 0.04 | 0.03 |
| *PF87* | 0.57 | 0.05 | 0.09 | 0.00 | 0.02 | 0.00 | 0.03 | 0.08 | -0.01 | 0.01 | 0.01 | 0.04 | -0.01 | 0.00 | 0.00 | -0.02 | -0.02 | 0.00 | 0.00 | 0.00 | -0.01 |
| *PF89* | 0.60 | 0.03 | 0.10 | 0.02 | 0.04 | 0.03 | 0.07 | 0.12 | 0.03 | 0.00 | 0.05 | 0.02 | -0.03 | 0.00 | 0.00 | 0.00 | 0.04 | 0.02 | 0.03 | 0.04 | 0.04 |
| *PF90* | 0.56 | 0.01 | 0.15 | -0.02 | 0.03 | -0.03 | 0.08 | 0.03 | 0.01 | 0.00 | 0.00 | 0.05 | 0.00 | -0.02 | 0.01 | 0.00 | -0.01 | -0.03 | -0.01 | -0.02 | -0.01 |
| *PF91* | 0.58 | 0.04 | 0.13 | -0.01 | 0.05 | 0.00 | 0.09 | 0.07 | 0.01 | 0.07 | 0.03 | 0.03 | 0.04 | -0.02 | 0.04 | -0.01 | 0.00 | 0.03 | 0.02 | -0.01 | -0.02 |
| *PF92* | 0.57 | 0.04 | 0.16 | -0.04 | 0.00 | 0.01 | 0.11 | 0.10 | 0.06 | 0.03 | 0.01 | 0.11 | -0.02 | 0.00 | 0.02 | -0.03 | 0.03 | 0.00 | 0.03 | 0.00 | 0.02 |
| *PF93* | 0.55 | 0.07 | 0.11 | 0.01 | 0.03 | -0.01 | 0.07 | 0.03 | 0.01 | 0.03 | -0.01 | 0.05 | 0.04 | 0.00 | 0.03 | -0.01 | 0.02 | 0.03 | 0.00 | -0.02 | -0.03 |
| *PF94* | 0.57 | 0.03 | 0.18 | -0.03 | -0.01 | -0.01 | 0.12 | 0.02 | 0.04 | 0.04 | -0.03 | 0.08 | 0.04 | 0.00 | 0.04 | -0.02 | -0.01 | 0.00 | -0.02 | 0.00 | -0.01 |
| *PF95* | 0.55 | 0.04 | 0.12 | 0.03 | 0.04 | 0.00 | 0.09 | 0.06 | 0.00 | 0.03 | 0.02 | 0.07 | 0.03 | -0.01 | 0.04 | -0.01 | -0.02 | 0.02 | -0.03 | -0.01 | 0.00 |

**Figure S2**: $F_{ST}$ genetic differentiation values between each pair of stations are visualized using a station- by- station matrix for Amitermes parvulus samples. Cells are colored according to the genetic differentiation between each pair of stations; darker shades of grey indicate that stations are more genetically similar to each other (i.e., more likely to belong to the same colony).
