## Supplemental Table S1 for "Comparative genetic study of the colony structure and colony spatial distribution between the higher termite *Amitermes parvulus* and the lower, subterranean termite *Reticulitermes flavipes* in an urban environment"

**Table S1**: Collection points sampled for the fine- and large-scale analyses. The genus and number of workers analyzed are indicated for each sample point, as well as its genetically-inferred colony of origin. The geographic coordinates are indicated for each sample point (position on the grid-site and GPS coordinates for, respectively fine- and large-scale analysis samples)

| Experiment | Sample # | Genus | Coordinates | | #workers analyzed | Genetically assigned Colony ID |
|---|---|---|---|---|---|---|
| | | | Column | Row | | |
| Fine-scale | *PF41* | *Amitermes* | 16 | 6 | 8 | A1 |
| Fine-scale | *PF54* | *Amitermes* | 2 | 2 | 5 | A2 |
| Fine-scale | *PF58* | *Amitermes* | 11 | 4 | 4 | A1 |
| Fine-scale | *PF59* | *Amitermes* | 12 | 4 | 5 | A1 |
| Fine-scale | *PF61* | *Amitermes* | 13 | 2 | 7 | A1 |
| Fine-scale | *PF63* | *Amitermes* | 13 | 5 | 8 | A1 |
| Fine-scale | *PF65* | *Amitermes* | 14 | 1 | 7 | A1 |
| Fine-scale | *PF68* | *Amitermes* | 14 | 4 | 7 | A1 |
| Fine-scale | *PF69* | *Amitermes* | 14 | 6 | 8 | A1 |
| Fine-scale | *PF70* | *Amitermes* | 14 | 7 | 8 | A1 |
| Fine-scale | *PF76* | *Amitermes* | 15 | 6 | 8 | A1 |
| Fine-scale | *PF79* | *Amitermes* | 15 | 7 | 8 | A1 |
| Fine-scale | *PF84* | *Amitermes* | 16 | 7 | 8 | A1 |
| Fine-scale | *PF87* | *Amitermes* | 17 | 4 | 8 | A1 |
| Fine-scale | *PF89* | *Amitermes* | 17 | 5 | 8 | A1 |
| Fine-scale | *PF90* | *Amitermes* | 17 | 6 | 8 | A1 |
| Fine-scale | *PF91* | *Amitermes* | 18 | 3 | 8 | A1 |
| Fine-scale | *PF92* | *Amitermes* | 18 | 5 | 8 | A1 |
| Fine-scale | *PF93* | *Amitermes* | 19 | 1 | 8 | A1 |
| Fine-scale | *PF94* | *Amitermes* | 19 | 3 | 8 | A1 |
| Fine-scale | *PF95* | *Amitermes* | 19 | 5 | 8 | A1 |
| Fine-scale | *PF1* | *Reticulitermes* | 1 | 1 | 10 | R1 |
| Fine-scale | *PF17* | *Reticulitermes* | 19 | 6.1 | 10 | R10 |
| Fine-scale | *PF52* | *Reticulitermes* | 1 | 7 | 10 | R11 |
| Fine-scale | *PF55* | *Reticulitermes* | 3 | 1 | 10 | R12 |
| Fine-scale | *PF56* | *Reticulitermes* | 5 | 4 | 10 | R13 |
| Fine-scale | *PF57* | *Reticulitermes* | 5 | 5 | 5 | R13 |
| Fine-scale | *PF97* | *Reticulitermes* | 19 | 7 | 10 | R14 |
| Fine-scale | *PF23* | *Reticulitermes* | 10 | 4 | 9 | R15 |
| Fine-scale | *PF26* | *Reticulitermes* | 12 | 5.2 | 10 | R16 |
| Fine-scale | *PF29* | *Reticulitermes* | 14 | 3 | 4 | R17 |
| Fine-scale | *PF66* | *Reticulitermes* | 14 | 3.2 | 10 | R17 |
| Fine-scale | *PF75* | *Reticulitermes* | 15 | 3 | 10 | R18 |
| Fine-scale | *PF81* | *Reticulitermes* | 16 | 1 | 10 | R19 |
| Fine-scale | *PF2* | *Reticulitermes* | 2 | 3 | 10 | R2 |
| Fine-scale | *PF3* | *Reticulitermes* | 3 | 3 | 10 | R2 |
| Fine-scale | *PF50* | *Reticulitermes* | 19 | 6.2 | 10 | R20 |
| Fine-scale | *PF51* | *Reticulitermes* | 19 | 6.3 | 10 | R20 |
| Fine-scale | *PF96* | *Reticulitermes* | 19 | 6 | 10 | R20 |
| Fine-scale | *PF53* | *Reticulitermes* | 1 | 7.1 | 10 | R21 |
| Fine-scale | *PF34* | *Reticulitermes* | 15 | 1 | 10 | R22 |
| Fine-scale | *PF73* | *Reticulitermes* | 15 | 2 | 10 | R22 |
| Fine-scale | *PF74* | *Reticulitermes* | 15 | 1.2 | 10 | R22 |
| Fine-scale | *PF77* | *Reticulitermes* | 15 | 6.2 | 10 | R23 |

| | | | | | | |
|---|---|---|---|---|---|---|
| Fine-scale | *PF82* | *Reticulitermes* | 16 | 2 | 10 | R24 |
| Fine-scale | *PF4* | *Reticulitermes* | 5 | 2 | 10 | R3 |
| Fine-scale | *PF5* | *Reticulitermes* | 10 | 4.4 | 10 | R4 |
| Fine-scale | *PF7* | *Reticulitermes* | 11 | 5 | 10 | R4 |
| Fine-scale | *PF8* | *Reticulitermes* | 12 | 5 | 10 | R5 |
| Fine-scale | *PF11* | *Reticulitermes* | 14 | 4 | 10 | R5 |
| Fine-scale | *PF67* | *Reticulitermes* | 14 | 4.3 | 10 | R5 |
| Fine-scale | *PF28* | *Reticulitermes* | 13 | 7 | 10 | R6 |
| Fine-scale | *PF14* | *Reticulitermes* | 16 | 5 | 11 | R7 |
| Fine-scale | *PF15* | *Reticulitermes* | 17 | 6 | 10 | R8 |
| Fine-scale | *PF16* | *Reticulitermes* | 19 | 5 | 10 | R9 |
| Large-scale | *Am_01* | *Amitermes* | -97.5821813 | 30.4735933 | 7 | A3 |
| Large-scale | *Am_02* | *Amitermes* | -97.5821739 | 30.4735806 | 7 | A3 |
| Large-scale | *Am_03* | *Amitermes* | -97.5821824 | 30.4731816 | 0 | / |
| Large-scale | *Am_04* | *Amitermes* | -97.5825518 | 30.4718570 | 6 | A4 |
| Large-scale | *Am_05* | *Amitermes* | -97.5826669 | 30.4713189 | 7 | A4 |
| Large-scale | *Am_06* | *Amitermes* | -97.5835050 | 30.4726870 | 0 | / |
| Large-scale | *Am_07* | *Amitermes* | -97.5853825 | 30.4722500 | 8 | A5 |
| Large-scale | *Am_08* | *Amitermes* | -97.5854623 | 30.4722084 | 8 | A5 |
| Large-scale | *Am_09* | *Amitermes* | -97.5860620 | 30.4716562 | 7 | A6 |
| Large-scale | *Am_10* | *Amitermes* | -97.5860980 | 30.4716804 | 7 | A6 |
| Large-scale | *Am_11* | *Amitermes* | -97.5861638 | 30.4717636 | 8 | A6 |
| Large-scale | *Am_12* | *Amitermes* | -97.5826560 | 30.4714770 | 6 | A4 |
| Large-scale | *Am_13* | *Amitermes* | -97.5825480 | 30.4714530 | 6 | A4 |
| Large-scale | *Am_14* | *Amitermes* | -97.5859157 | 30.4722416 | 7 | A5 |
| Large-scale | *Am_15* | *Amitermes* | -97.5857695 | 30.4723768 | 7 | A5 |
| Large-scale | *Am_16* | *Amitermes* | -97.5853112 | 30.4731674 | 7 | A7 |
| Large-scale | *Am_17* | *Amitermes* | -97.5852857 | 30.4732749 | 7 | A7 |
| Large-scale | *Am_18* | *Amitermes* | -97.5857790 | 30.4747414 | 4 | A8 |
| Large-scale | *Am_19* | *Amitermes* | -97.5856820 | 30.4746713 | 5 | A9 |
| Large-scale | *Am_20* | *Amitermes* | -97.5865287 | 30.4750904 | 4 | A10 |
| Large-scale | *Am_21* | *Amitermes* | -97.5868507 | 30.4751346 | 6 | A11 |
| Large-scale | *Am_22* | *Amitermes* | -97.5869057 | 30.4751392 | 7 | A12 |
| Large-scale | *Am_23* | *Amitermes* | -97.5882900 | 30.4754800 | 7 | A13 |
| Large-scale | *Am_24* | *Amitermes* | -97.5883000 | 30.4752400 | 0 | / |
| Large-scale | *Am_25* | *Amitermes* | -97.5882150 | 30.4717240 | 5 | A14 |
| Large-scale | *Am_26* | *Amitermes* | -97.5897470 | 30.4715880 | 0 | / |
| Large-scale | Retic_MM1 | *Reticulitermes* | -97.5907368 | 30.4721323 | / | / |
| Large-scale | Retic_KL1 | *Reticulitermes* | -97.5890218 | 30.4710160 | / | / |
| Large-scale | Retic_MM10 | *Reticulitermes* | -97.5898444 | 30.4716921 | / | / |
| Large-scale | Retic_MM11 | *Reticulitermes* | -97.5897706 | 30.4717291 | / | / |
| Large-scale | Retic_MM12 | *Reticulitermes* | -97.5897304 | 30.4715870 | / | / |
| Large-scale | Retic_MM13 | *Reticulitermes* | -97.5892718 | 30.4716656 | / | / |
| Large-scale | Retic_MM14 | *Reticulitermes* | -97.5887554 | 30.4717222 | / | / |
| Large-scale | Retic_MM15 | *Reticulitermes* | -97.5886495 | 30.4716690 | / | / |
| Large-scale | Retic_MM16 | *Reticulitermes* | -97.5884805 | 30.4716737 | / | / |
| Large-scale | Retic_MM17 | *Reticulitermes* | -97.5881975 | 30.4717569 | / | / |
| Large-scale | Retic_MM18 | *Reticulitermes* | -97.5880983 | 30.4716933 | / | / |
| Large-scale | Retic_MM19 | *Reticulitermes* | -97.5879897 | 30.4717268 | / | / |
| Large-scale | Retic_MM2 | *Reticulitermes* | -97.5906603 | 30.4720965 | / | / |
| Large-scale | Retic_MM20 | *Reticulitermes* | -97.5893239 | 30.4713003 | / | / |
| Large-scale | Retic_MM21 | *Reticulitermes* | -97.5885085 | 30.4714760 | / | / |

| | | | | | | |
|---|---|---|---|---|---|---|
| Large-scale | Retic_MM22 | *Reticulitermes* | -97.5887285 | 30.4713974 | / | / |
| Large-scale | Retic_MM23 | *Reticulitermes* | -97.5884495 | 30.4713419 | / | / |
| Large-scale | Retic_MM24 | *Reticulitermes* | -97.5861327 | 30.4711680 | / | / |
| Large-scale | Retic_MM25 | *Reticulitermes* | -97.5861380 | 30.4711044 | / | / |
| Large-scale | Retic_MM26 | *Reticulitermes* | -97.5859154 | 30.4711576 | / | / |
| Large-scale | Retic_MM27 | *Reticulitermes* | -97.5858215 | 30.4713009 | / | / |
| Large-scale | Retic_MM28 | *Reticulitermes* | -97.5878926 | 30.4716231 | / | / |
| Large-scale | Retic_MM29 | *Reticulitermes* | -97.5875654 | 30.4714000 | / | / |
| Large-scale | Retic_MM3 | *Reticulitermes* | -97.5905222 | 30.4720329 | / | / |
| Large-scale | Retic_MM30 | *Reticulitermes* | -97.5877115 | 30.4711376 | / | / |
| Large-scale | Retic_MM31 | *Reticulitermes* | -97.5880012 | 30.4709076 | / | / |
| Large-scale | Retic_MM32 | *Reticulitermes* | -97.5880039 | 30.4707816 | / | / |
| Large-scale | Retic_MM33 | *Reticulitermes* | -97.5883392 | 30.4708602 | / | / |
| Large-scale | Retic_MM34 | *Reticulitermes* | -97.5884277 | 30.4706891 | / | / |
| Large-scale | Retic_MM35 | *Reticulitermes* | -97.5885471 | 30.4708498 | / | / |
| Large-scale | Retic_MM36 | *Reticulitermes* | -97.5886369 | 30.4706198 | / | / |
| Large-scale | Retic_MM37 | *Reticulitermes* | -97.5889105 | 30.4708382 | / | / |
| Large-scale | Retic_MM38 | *Reticulitermes* | -97.5891184 | 30.4708244 | / | / |
| Large-scale | Retic_MM39 | *Reticulitermes* | -97.5892645 | 30.4707319 | / | / |
| Large-scale | Retic_MM4 | *Reticulitermes* | -97.5904176 | 30.4719902 | / | / |
| Large-scale | Retic_MM40 | *Reticulitermes* | -97.5890231 | 30.4705816 | / | / |
| Large-scale | Retic_MM41 | *Reticulitermes* | -97.5896521 | 30.4704996 | / | / |
| Large-scale | Retic_MM42 | *Reticulitermes* | -97.5891881 | 30.4706383 | / | / |
| Large-scale | Retic_MM43 | *Reticulitermes* | -97.5877487 | 30.4708140 | / | / |
| Large-scale | Retic_MM44 | *Reticulitermes* | -97.5871251 | 30.4712509 | / | / |
| Large-scale | Retic_MM45 | *Reticulitermes* | -97.5868917 | 30.4714335 | / | / |
| Large-scale | Retic_MM46 | *Reticulitermes* | -97.5865940 | 30.4715976 | / | / |
| Large-scale | Retic_MM47 | *Reticulitermes* | -97.5819305 | 30.4734771 | / | / |
| Large-scale | Retic_MM48 | *Reticulitermes* | -97.5836203 | 30.4743509 | / | / |
| Large-scale | Retic_MM49 | *Reticulitermes* | -97.5833038 | 30.4750976 | / | / |
| Large-scale | Retic_MM50 | *Reticulitermes* | -97.5835090 | 30.4762811 | / | / |
| Large-scale | Retic_MM51 | *Reticulitermes* | -97.5834298 | 30.4763909 | / | / |
| Large-scale | Retic_MM52 | *Reticulitermes* | -97.5834620 | 30.4765504 | / | / |
| Large-scale | Retic_MM53 | *Reticulitermes* | -97.5834915 | 30.4764672 | / | / |
| Large-scale | Retic_MM54 | *Reticulitermes* | -97.5836082 | 30.4763089 | / | / |
| Large-scale | Retic_MM55 | *Reticulitermes* | -97.5837396 | 30.4762915 | / | / |
| Large-scale | Retic_MM56 | *Reticulitermes* | -97.5838375 | 30.4763447 | / | / |
| Large-scale | Retic_MM57 | *Reticulitermes* | -97.5835975 | 30.4765528 | / | / |
| Large-scale | Retic_MM58 | *Reticulitermes* | -97.5837598 | 30.4764753 | / | / |
| Large-scale | Retic_MM59 | *Reticulitermes* | -97.5837034 | 30.4766071 | / | / |
| Large-scale | Retic_MM6 | *Reticulitermes* | -97.5903747 | 30.4719647 | / | / |
| Large-scale | Retic_MM60 | *Reticulitermes* | -97.5838268 | 30.4766637 | / | / |
| Large-scale | Retic_MM61 | *Reticulitermes* | -97.5839019 | 30.4765724 | / | / |
| Large-scale | Retic_MM62 | *Reticulitermes* | -97.5839220 | 30.4764568 | / | / |
| Large-scale | Retic_MM63 | *Reticulitermes* | -97.5837343 | 30.4768822 | / | / |
| Large-scale | Retic_MM64 | *Reticulitermes* | -97.5837705 | 30.4767712 | / | / |
| Large-scale | Retic_MM65 | *Reticulitermes* | -97.5833896 | 30.4768660 | / | / |
| Large-scale | Retic_MM66 | *Reticulitermes* | -97.5835573 | 30.4770578 | / | / |
| Large-scale | Retic_MM67 | *Reticulitermes* | -97.5834294 | 30.4771809 | / | / |
| Large-scale | Retic_MM68 | *Reticulitermes* | -97.5834508 | 30.4781194 | / | / |
| Large-scale | Retic_MM69 | *Reticulitermes* | -97.5832389 | 30.4775184 | / | / |
| Large-scale | Retic_MM7 | *Reticulitermes* | -97.5902727 | 30.4719278 | / | / |
| Large-scale | Retic_MM8 | *Reticulitermes* | -97.5900729 | 30.4718515 | / | / |

| | | | | | | |
|---|---|---|---|---|---|---|
| Large-scale | Retic_MM9 | *Reticulitermes* | -97.5898484 | 30.4716124 | / | / |
| Large-scale | Retic_PA1 | *Reticulitermes* | -97.5869037 | 30.4716813 | / | / |
| Large-scale | Retic_PA10 | *Reticulitermes* | -97.5856810 | 30.4721583 | / | / |
| Large-scale | Retic_PA11 | *Reticulitermes* | -97.5853863 | 30.4728981 | / | / |
| Large-scale | Retic_PA12 | *Reticulitermes* | -97.5853138 | 30.4733628 | / | / |
| Large-scale | Retic_PA14 | *Reticulitermes* | -97.5852924 | 30.4735697 | / | / |
| Large-scale | Retic_PA15 | *Reticulitermes* | -97.5853004 | 30.4736367 | / | / |
| Large-scale | Retic_PA16 | *Reticulitermes* | -97.5853205 | 30.4737708 | / | / |
| Large-scale | Retic_PA17 | *Reticulitermes* | -97.5851784 | 30.4739696 | / | / |
| Large-scale | Retic_PA18 | *Reticulitermes* | -97.5852012 | 30.4740563 | / | / |
| Large-scale | Retic_PA19 | *Reticulitermes* | -97.5854142 | 30.4744039 | / | / |
| Large-scale | Retic_PA2 | *Reticulitermes* | -97.5868822 | 30.4717692 | / | / |
| Large-scale | Retic_PA20 | *Reticulitermes* | -97.5854370 | 30.4744617 | / | / |
| Large-scale | Retic_PA21 | *Reticulitermes* | -97.5855135 | 30.4745426 | / | / |
| Large-scale | Retic_PA22 | *Reticulitermes* | -97.5855577 | 30.4745749 | / | / |
| Large-scale | Retic_PA23 | *Reticulitermes* | -97.5856060 | 30.4746119 | / | / |
| Large-scale | Retic_PA24 | *Reticulitermes* | -97.5860901 | 30.4749517 | / | / |
| Large-scale | Retic_PA25 | *Reticulitermes* | -97.5861827 | 30.4749968 | / | / |
| Large-scale | Retic_PA26 | *Reticulitermes* | -97.5862591 | 30.4750188 | / | / |
| Large-scale | Retic_PA27 | *Reticulitermes* | -97.5863181 | 30.4750511 | / | / |
| Large-scale | Retic_PA28 | *Reticulitermes* | -97.5863865 | 30.4750592 | / | / |
| Large-scale | Retic_PA29 | *Reticulitermes* | -97.5864710 | 30.4750812 | / | / |
| Large-scale | Retic_PA3 | *Reticulitermes* | -97.5868179 | 30.4717333 | / | / |
| Large-scale | Retic_PA30 | *Reticulitermes* | -97.5867287 | 30.4751230 | / | / |
| Large-scale | Retic_PA4 | *Reticulitermes* | -97.5867602 | 30.4717414 | / | / |
| Large-scale | Retic_PA5 | *Reticulitermes* | -97.5866381 | 30.4718200 | / | / |
| Large-scale | Retic_PA6 | *Reticulitermes* | -97.5865604 | 30.4719009 | / | / |
| Large-scale | Retic_PA7 | *Reticulitermes* | -97.5864571 | 30.4719460 | / | / |
| Large-scale | Retic_PA8 | *Reticulitermes* | -97.5864008 | 30.4719472 | / | / |
| Large-scale | Retic_PA9 | *Reticulitermes* | -97.5862036 | 30.4720581 | / | / |
